# Peroxisome biogenesis initiated by protein phase separation

**DOI:** 10.1101/2022.09.20.508718

**Authors:** Rini Ravindran, Isabel O. L. Bacellar, Xavier Castellanos-Girouard, Zhenghao Zhang, Lydia Kisley, Stephen W. Michnick

**Author notes:** IOLB’s present address is Douglas Research Centre, 6875 LaSalle Blvd., Montreal, Quebec, H4H 1R3, Canada. ZZ’s present address is Mitchell Physics Building (MPHY); 578 University Drive; College Station, Texas 77843-4242, USA.

## Abstract

Peroxisomes are organelles that perform beta-oxidation of fatty acids and amino acids. Both rare and prevalent diseases are caused by their disfunction^1^. Among disease-causing mutant genes are those required for protein transport into the peroxisome. The peroxisomal protein import machinery, also shared with chloroplasts, is unique in transporting folded and large, up to 10 nm in diameter, protein complexes into peroxisomes^2^ and current models postulate a large pore formed by transmembrane proteins^3^. To date, however, no pore structure has been observed. In the budding yeast *Saccharomyces cerevisiae*, the minimum transport machinery includes membrane proteins Pex13 and Pex14 and cargo protein-binding transport receptor, Pex5. Here we show that Pex13 undergoes liquid-liquid phase separation (LLPS) with Pex5-cargo. Intrinsically disordered regions (IDR) in Pex13 and Pex5 resemble those found in nuclear pore complex (NPC) proteins. Cargo transport into peroxisomes depends on the number but not patterns of aromatic residues in these IDRs, consistent with their roles as ‘stickers’ in associative polymer models of LLPS^4,5^. Finally, imaging Fluorescence Cross-Correlation Spectroscopy (iFCCS) shows that the transport of cargo correlates with transient focusing of GFP-Pex13/14 on the peroxisome membrane. Pex13 and Pex14 form foci in distinct time-frames, suggesting that they may form channels at different saturating concentrations of Pex5-cargo. Our results suggest a model in which LLPS of Pex5-cargo with Pex13/14 results in transient protein transport channels.

## Main Text

The peroxins Pex5, Pex13, and Pex14 play a critical role in the protein translocation step of the peroxisome transmembrane transport PTS1 pathway. Pex5 binds to cargo proteins in the cytosol, by recognition of peroxisomal targeting signal (PTS1) motifs – the C-terminal tripeptide sequence serine-lysine-leucine (-SKL or its variants)^6^. A conformation change upon binding cargo exposes an IDR, which interacts with the peroxisome membrane proteins Pex13 and Pex14, leading to translocation of cargo proteins to the peroxisomal matrix^7^. No channel structure has ever been observed in peroxisomes and no physical model of transport has been validated but compelling evidence of at least transiently formed channels has been provided. An *in vitro* study where lipid membranes reconstituted with Pex14 complexes showed Pex5-cargo-dependent ion conductance with cargos of up to 9 nm in diameter^3^.

A clue to a possible mechanism of transport is the sequence characteristics of IDRs in Pex5, Pex13 and Pex14 that resemble those of nucleoporin proteins of nuclear pore complexes (Fig. 1a,b). Notably, Pex13 features a series of Tyr-Gly (YG) repeats within its IDR (Fig. 1c). We hypothesized that these residues may undergo LLPS by engaging in π-π or π-cation interactions similar to those formed by Phe-Gly (FG) repeats in nucleoporins that undergo LLPS and compose the selective-permeability filter of the NPC; we postulate that the resulting condensate forms a simple, primitive, and transient NPC-like assembly^8,9^. This hypothesis would provide a natural explanation for protein import to peroxisomes, with peroxins forming a biomolecular condensate through LLPS and creating a conduit for cargo transport across the peroxisomal membrane, with Pex5 acting as a shuttling receptor analogously to karyopherins^10^. The budding yeast *Saccharomyces cerevisiae* Pex13 has aromatic repeats in its IDR and Pex5 and Pex13 notably have prion-like domains (PLD) within their IDRs, which have been demonstrated to form biomolecular condensates through multivalent π-π or π-cation ‘sticker’ interactions^4,5,11–13^. Indeed, peroxisomal proteins were shown to precipitate as large complexes under ATP depletion, suggesting that their formation precede translocation^14,15^. We thus set out to test the hypothesis that peroxisome protein transport occurs through Pex5-cargo-Pex13-Pex14 LLPS.

**Fig. 1.**
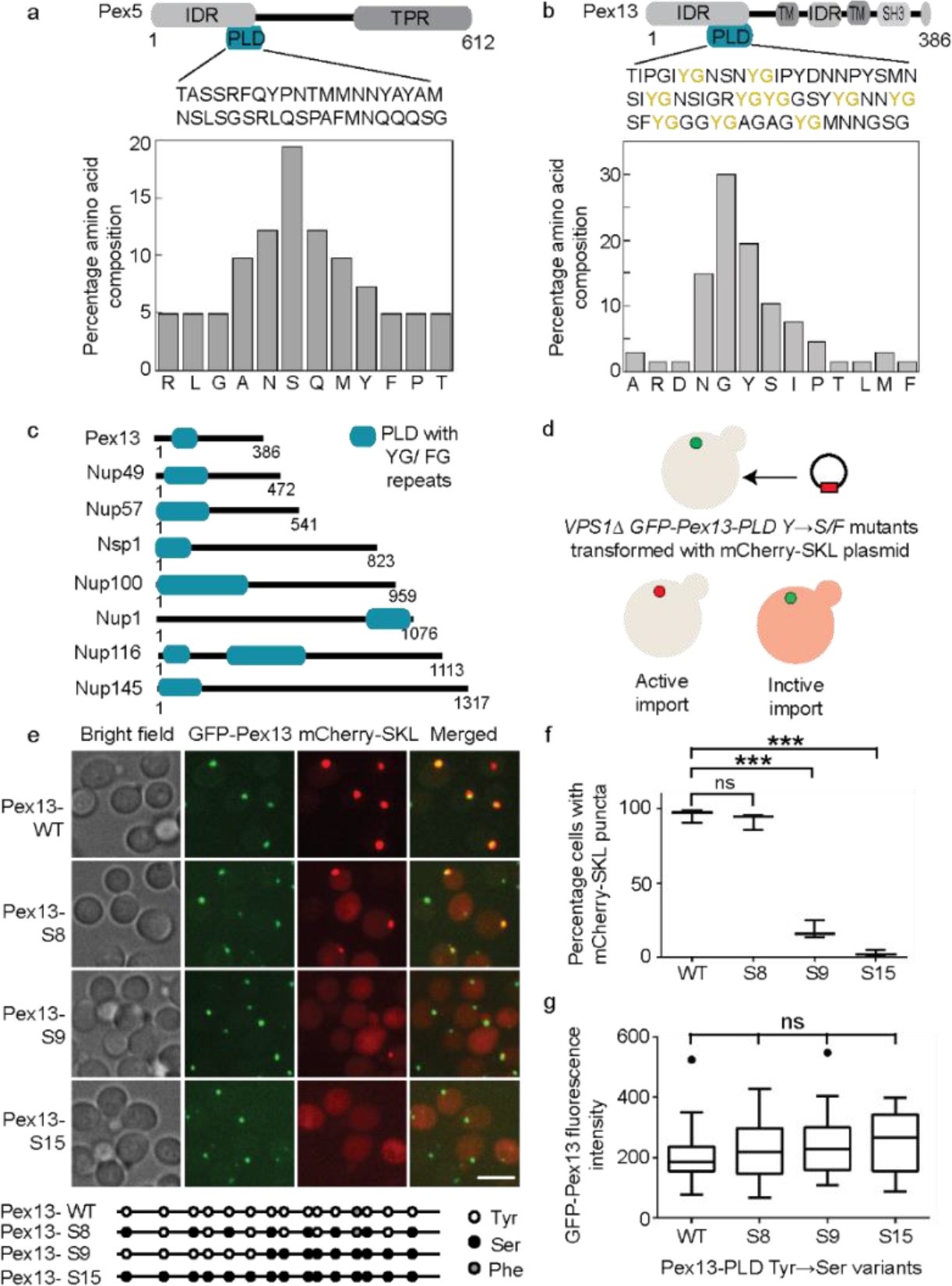
Pex13 Tyr to Ser mutations disrupt peroxisomal PTS1-cargo import. **a**, Domain structures of Pex5 and **b**, Pex13, highlighting intrinsically disordered domains (IDR), tetratricopeptide repeat (TPR), transmembrane domains (TM) and SRC homology 3 domain (SH3). Percentage amino acid composition of the prion-like domains (PLD) within the IDR is presented below the PLD sequence. **c**, FG -rich PLD regions within Pex13 and the nucleoporin proteins that line the central channel of the nuclear pore complex. **d**, Schematic diagram of the assay to measure PTS1 import. *VPS1*Δ deletion strain expressing GFP-Pex13 with or without Tyr to Ser or Phe mutations in the Pex13 PLD were transformed with mCherry-SKL expression plasmid. Cells with active PTS1 import show red mCherry-SKL puncta that overlap with GFP-Pex13, while mCherry-SKL remains cytoplasmic in cells with inactive PTS1 import. **e**, Confocal microscopy images of the indicated strains expressing mCherry-SKL. Scale bar 5 μM. Schematic of the Tyr to Ser mutations in Pex13-PLD is given below panel e. **f**, Percentage cells displaying mCherry-SKL foci imaged in panel e. Data are mean ± s.e.m. of n = 3 for 3 biologically independent experiments. \*\*\**P* < 0.001, ns *P*>0.05 one-way ANOVA with Tukey’s multiple comparison test. **g**, GFP-Pex13 fluorescence intensity in the indicated strains. n=50 cells each for 3 biologically independent experiments. ns *P*>0.05 one-way ANOVA Kruskal-Wali’s test with Dunn’s multiple comparison. The box limits represent the range between the first and third quartiles for each condition, the center lines show the median, and the ends of the whiskers extend to 1.5× the interquartile range.

## Results

### Peroxisomal protein transport depends on the number of Tyr-Gly repeats in Pex13

We predicted IDRs of yeast PTS1 peroxins, Pex5^16^, Pex13 and Pex14^12^, including PLD, within the N-terminal IDR regions of Pex5 and Pex13, but not in Pex14. (Extended data Fig. 1)^17^. The Pex5 PLD (residues 150-190) is enriched in polar amino acids, Ser (S), Gln (Q) and Asn (N) (Fig. 1a) a hallmark of several known prion-like proteins^11^. In contrast, the Pex13 PLD (amino acids 76-142) contains a high proportion of Tyr (Y) and Gly (G) residues forming a YG repeat domain, punctuated by S/N residues (Fig. 1b), reminiscent of Phe-Gly (FG) motifs with S/T spacers found in nucleoporin proteins that line the central channel of the NPC (Fig. 1c).

The Tyr residues within the Pex13 PLD are conserved despite very low sequence conservation (Extended Data Fig. 2b). To determine whether the YG repeats in Pex13 are important for PTS1 protein import, we used CRISPR-Cas9 to generate mutants in which Pex13-PLD YG motifs were replaced by SG or FG motifs. The mutations (S3-S15 and F1-F14) were made between Y71 and Y133, varying in number, pattern (blocky or interspersed), and location (Extended Data Fig. 2a). While the Y to S mutation preserves the hydroxyl group that engages in dipolar and hydrogen bond interactions, the FG change retains the aromatic ring required for π- π or π-cation interactions. To investigate whether the peroxisomal protein import is functional in the Pex13 YG mutants, we constitutively expressed mCherry fused to a PTS1-targeting SKL sequence as a model cargo protein for which we could visualize and quantitate peroxisomal uptake *in vivo* (Fig. 1d).

We observed that both S and F substitutions affect the transport of mCherry-SKL. While mutants with less than seven Y to S/F changes displayed mCherry-SKL foci like *WT* cells, mutants with more than 7 substitutions (*S7-S9 and F7-F9*) showed fewer mCherry-SKL foci (Extended Data Fig. 2a). No mCherry-SKL foci were observed for the S13-S15 and F13-F14 mutants showing that the Tyr residues in the Pex13 PLD are essential for PTS1 protein transport. To facilitate imaging experiments, we repeated the same assay in Pex13 YG mutants in which the *VPS1* gene is deleted, resulting in enlarged and mostly single peroxisomes in individual cells^18,19^. Pex13 was endogenously tagged to GFP, serving as a peroxisomal marker. We found that more than 90% of the *Pex13-WT* and *Pex13-S8* mutant cells displayed a discrete mCherry-SKL focus that overlaps with Pex13-GFP (Fig, 1e,f), thus showing active peroxisomal transport. In contrast, the frequency of cells with foci was reduced in Pex13-S9 mutants, with less than 25% of the S9 mutant cells showing mCherry-SKL foci. mCherry-SKL fluorescence remained cytoplasmic in cells expressing the Pex13-S15 variant lacking all aromatic amino acids within the PLD (Fig, 1e-g). The sharp transition in mCherry-SKL import at a critical number of Tyr residues is consistent with observed dependencies of LLPS on numbers of aromatic *π*-*π* or *π*-cation interactions that can be made^4^.

### Number but not pattern of YG repeats in Pex13 determine protein transport efficiency

To further investigate the role of Pex13 YG repeats in peroxisomal protein import, we designed an assay to monitor mCherry-SKL import as a function of time, and therefore mCherry-Pex5 complex abundance, by transiently expressing the *PEX5* PTS1-specific transporter gene. We introduced the inducible *pGAL1* promoter immediately upstream of the *PEX5* gene, thus allowing for Pex5 expression and transport of PTS1 cargo only when cells are grown in galactose^20^. We measured the kinetics of mCherry-SKL foci formation in Pex13-WT and Pex13 Y to S mutants expressing pGal1-Pex5 by switching cells from a raffinose to galactose-containing medium (Fig. 2a). We found that the transport of mCherry-SKL begins 30 min after Pex5 induction in WT cells, with over 90% of cells showing mCherry foci within 1 hour (Fig. 2b,c). mCherry-SKL foci appear in an all-or-none manner that we can resolve as happening over less than one minute, followed by a continuous increase in intensity (Fig. 2d), suggesting that Pex5 expression must reach a critical level for protein import to take place. Similarly, the S8 and S9 mutant strains showed all-or-none appearance of foci, but only after 120 and 240 minutes of Pex5 induction, respectively (Fig. 2e, Extended Data Fig. 3). These results imply that saturating concentrations of Pex5 for LLPS increase with the number of Tyr to Ser substitutions and rates of uptake decrease.

**Fig. 2.**
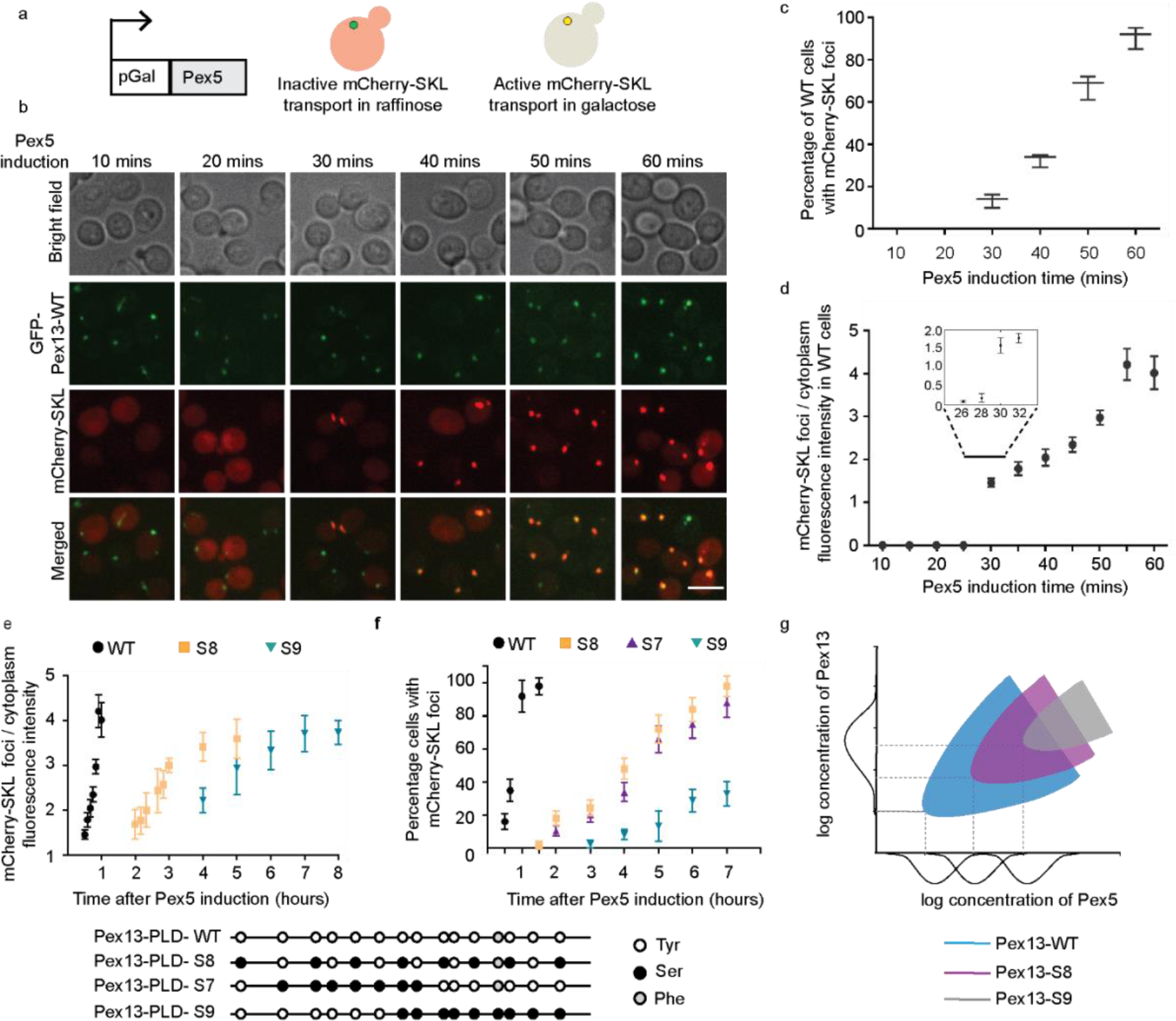
Pex13-PLD Tyr to Ser mutations increase threshold Pex5 concentration and decrease rates of PTS1-cargo transport. **a**, Phenotype of cells grown in raffinose (repression of Pex5 expression) and galactose (induction of Pex5 expression). mCherry-SKL cargo will only be transported into peroxisomes if it is bound to Pex5. (*left*) mCherry-SKL remains cytoplasmic in cells grown in raffinose (no Pex5 expression) but (*right*), mCherry-SKL puncta overlaps with GFP-Pex13 following galactose induction of Pex5 expression. **b**, Spinning disc confocal images of Pex13-GFP WT cells after induction of Pex5. Scale bar 5 μM. **c**, Percentage of cells displaying mCherry-SKL foci imaged in panel **b**. Error bars are standard errors of mean for three independent experiments. n=50 cells. **d**, Cargo transport kinetics measured as the ratio of pixel sum intensity of mCherry-SKL foci superimposed on Pex13-GFP to the cytoplasmic mCherry-SKL fluorescence measured in GFP-Pex13-WT cells following induction of Pex5 expression. The inset indicates all-or-none appearance of mCherry-SKL puncta at 2 minutes resolution. **e**, (*lower*) mCherry-SKL cargo transport kinetics and **f**, Percentage of cells displaying mCherry-SKL foci in the indicated strains following Pex5 induction. Error bars are standard errors of mean for three independent experiments, n=50 cells. **e**, (*upper*) Schematic representation of the sequence distributions of Tyr to Ser mutations in the Pex13 PLD mutants indicated above. **g**, Schematic representation of hypothetical effects of Tyr to Ser mutations on phase separation of Pex5-Pex13 complexes. Phase separation of Pex13 WT occurs when expression of Pex5 reaches a saturation concentration at which it is above the phase boundary for phase separation (blue U-shaped boundary). Increasing numbers of Tyr to Ser mutations in Pex13 result in right- and upward-shifts in the phase boundary with corresponding to increases in the saturating concentration of Pex5. Distributions to the left or under the y and x axes, respectively, represent distributions of Pex13 and Pex5 abundances across a population of cells.

In our hypothesis, these observations are consistent with transport following phase separation that occurs when Pex5 reaches its saturating concentration for LLPS with Pex13. Both saturating concentration and rate of protein transport depend on the strengths of intermolecular aromatic *π*-*π* or *π*-cation interactions that can be made between Pex13, Pex14, and Pex5 IDRs. Decreased valency of the S7-9 mutants result in the reduction of favorable free energies of phase separation^4^. Furthermore, the strengths of interactions and patterns of YG repeats are important when we consider the frequency of mCherry-SKL foci in a population of cells (Fig. 2f). Here we observe that blocky or interspersed Tyr to Ser mutations of similar number (S7 or S8, respectively) have the same effect on frequency of mCherry-SKL foci, whereas just one more Tyr to Ser mutation (S9) reduces the frequency much further, and steady-state frequencies of foci are reduced significantly (Fig. 1f, Extended Data Fig. 4,5). These results follow from expected variations in expression of Pex13/14 and Pex5 across a population of cells and how this could determine saturating concentrations of Pex5 if we project concentration variations onto an LLPS phase diagram (Fig. 2g)^21^. Pex5 and Pex13 mean abundances and variations are almost equal and assuming a Gaussian distribution of abundances would vary from approximately 1,000 to 2,000 copies per cell with mean of about 1,600 copies. Assuming saturating concentrations of Pex5 and Pex13 are equal to the mean concentrations of the protein, then we would expect to observe LLPS of Pex5-cargo/Pex13/Pex14 and consequently cargo uptake in most to all cells. A small shift in the phase diagram for the S8 mutant, and consequently an increase in the saturating concentration, would only slightly reduce the number of cells where LLPS could occur, but a larger shift for S9 corresponding to about one standard deviation would lead to a reduction to about 20% of cells where protein transport will occur, as was observed (Fig. 2e).

### mCherry SKL cargo partitioning into Pex13-IDR condensates is Pex5-dependent *in vitro*

We expressed and purified the IDR regions of Pex13-WT, Pex13-S8, Pex13-S15 and Pex14 (Extended data Fig. 6) and induced LLPS by rapidly diluting the protein solution in a phase separation buffer similar to that used to study LLPS with nucleoporin IDRs (20 mM Tris pH 7.4 containing 10% PEG 8000)^9^. This assay confirmed that Pex13-IDR-WT undergoes salt and PEG-dependent phase separation (Extended Data Fig. 7a,b) forming spherical condensates at 10 μM, Pex13-IDR-S8 at 30 μM, and Pex13-IDR-S15 does not form condensates up to 100 μM (Fig. 3a), consistent with our hypothesis that saturating concentrations for condensation of peroxins depends on aromatic *π*-*π* or *π*-cation interactions. Pex13-IDR condensates readily fuse into single larger spheres with an average fusion timescale (⊤ _avg_) of 0.2 seconds (Fig. 3b and Extended Data Video1,2). Pex5 IDR also undergoes salt and PEG-dependent phase separates at 2 μM concentration (Extended Data Fig. 7c,d) In contrast, Pex14 IDR (4μM) forms elongated particles dependent on crowding but independent of salt concentration (Extended Data Fig. 7e,f).

**Fig 3.**
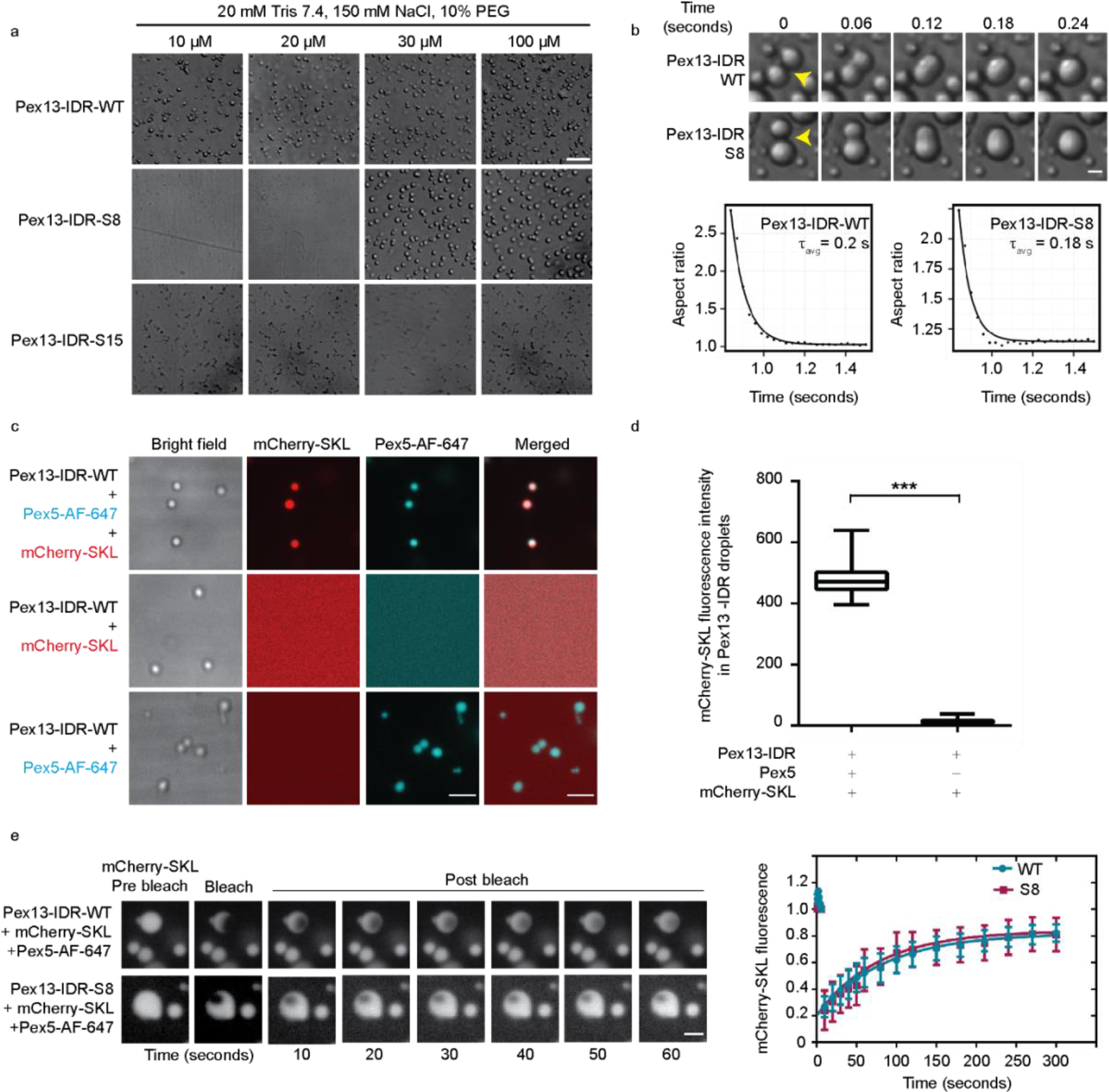
mCherry-SKL requires binding to Pex5 to partition into Pex13-IDR condensates *in vitro*. **a**, Bright field confocal images of condensates formed by purified Pex13-IDR-WT, S8 and S15 protein at the indicated concentrations at room temperature. Scale bar 50 μm. **b**, Fusion of Pex13-ISR and S8 purified protein in buffer containing 25 mM Tris, 150mM NaCl and 10% PEG at the indicated time points. Scale bar 5 μm. Fusion events are well fit by an exponential decay, which is used to determine fusion time-constant ⊤. The ⊤ _avg_ was calculated with data for n=18 condensates. **c**, Reconstitution experiments to measure partitioning of Pex5-AF 647 and mCherry-SKL into Pex13-IDR-WT condensates immediately after mixing at room temperature. Scale bar 5 μm. **d**, Fluorescence intensity of mCherry-SKL partitioned into Pex13-IDR condensates. n=30 condensates each from three independent experiments. \*\*\**P*<0.001 two-tailed Student’s t-test. The box limits represent the range between the first and third quartiles for each condition, the center lines show the median, and the ends of the whiskers extend to 1.5× the interquartile range. **e**, Representative images (left) of mCherry-SKL partitioned into Pex13-IDR-WT/S8 in the presence of Pex5 before and after photobleaching. FRAP quantification (right) of n=5 condensates imaged for 300 seconds post-bleaching. One-phase exponential equations were fitted to the curves. The average fluorescence before photobleaching was counted as 100%. Scale bar 2 μm.

We next tested whether mCherry-SKL partitioning into Pex13 IDR condensates is Pex5-dependent as we would expect for channel-forming events *in vivo*. We tagged purified recombinant full length Pex5 with AF dye 647 and titrated the concentrations at which Pex5 and mCherry-SKL do not form condensates in the phase separation buffer (Extended Fig. 8a-c). Under these conditions, Pex5 was incubated with mCherry-SKL followed by introduction of Pex13-IDR. Pex5-mCherry-SKL complexes partitioned into the Pex13-IDR-WT condensates (Fig. 3c,d, Extended Fig. 8d), while mCherry-SKL alone did not. Pex5 alone could also partition into Pex13 IDR condensates. Pex5-mCherry-SKL also partitioned into Pex13-IDR-S8 condensates at an elevated saturation concentration of 30 μM, and Pex13-IDR-S15 did not form condensates (Extended Fig. 8e). Thus, LLPS of Pex13 is required for partitioning of Pex5-cargo into Pex13 condensates *in vitro*. Further, mCherry-SKL partitioned within Pex13-IDR condensates in the presence of Pex5 shows a FRAP recovery within 2 minutes, but with a residual of 20%, suggesting a viscoelastic structure (Fig. 3e).

### Pex13 and Pex14 may form transient channels in peroxisome membrane

A model for peroxin-dependent protein transport begins to emerge by combining our evidence for Pex5-dependent partitioning of cargo into Pex13 condensates with that of transiently formed Pex5-cargo-dependent ion conductance in lipid bilayer vesicles reconstituted with Pex14 complexes^3^. It remains, however, that no transport pores in peroxisomes have been observed. We reason that these channels could be too small and too short lived to be directly observable with optical or electron tomographic techniques – note, for instance, that yeast peroxisomes on their own are already smaller than the diffraction limit. We instead sought to simply identify spatially-defined and transient correlations of cargo and peroxins as evidence of channel formation using imaging fluorescence cross correlation spectroscopy (iFCCS)^22,23^.

iFCCS calculates where mCherry-SKL and Pex13-GFP intensities are similar within individual peroxisomes and how the relative intensities change over milliseconds to seconds time scales (Extended Data Fig. 9). Microscopic images of Pex13-GFP and mCherry-SKL channels (Fig. 4a) were aligned using fiduciary markers and peroxisomes were identified using a previously established single particle tracking algorithm^24^. The intensities of Pex13-GFP or GFP-Pex14 and mCherry-SKL at each individual pixel within the peroxisome were cross-correlated over time. A cutoff of the magnitude of the cross-correlation amplitude, G_XC_(⊤) at ⊤ = 0 ms (i.e., the first frame) of ≥ 0.5 was selected to indicate peroxisomes of interest that agree with the prior mCherry-SKL foci data (Fig. 4j). Both spatial and temporal information could then be obtained by inspecting the correlation curves at each pixel and over the lag times (Extended Data Fig. 9f, g). We note that the emission of the fluorescent proteins is diffraction-limited, resulting in a ~300 nm resolution compared to the ~1000 nm × 100 nm VPS1 knockout strain peroxisomes. This low resolution does not matter because our goal was to capture spatiotemporally-resolved correlations of fluorescence sufficient to define single events, not to visualize channels, which, we reasoned, would likely be too dynamic to resolve with localization-based super-resolution methods.

**Fig. 4.**
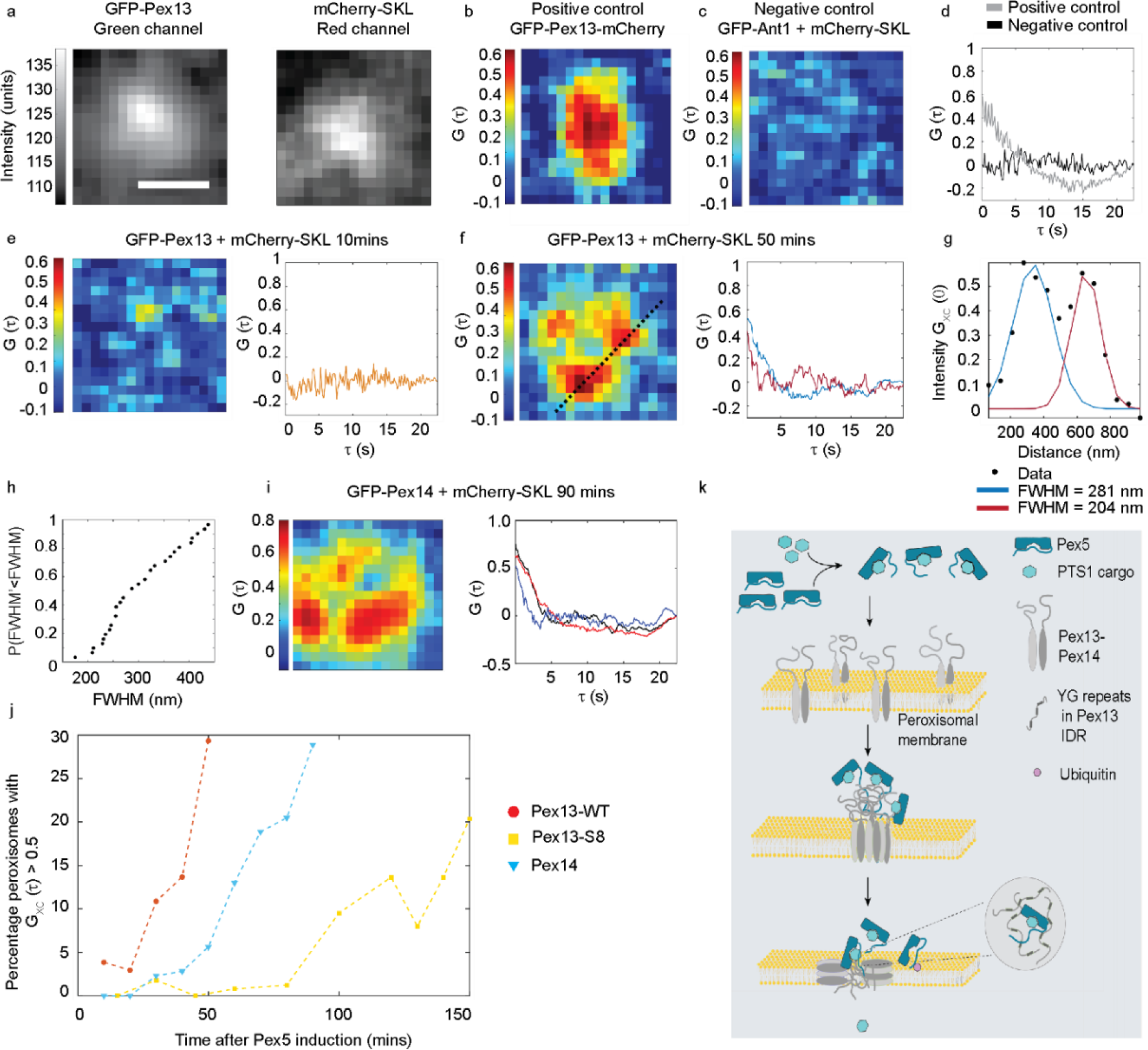
Multiple clusters of transiently correlated mCherry-SKL and Pex13-GFP in individual peroxisomes. **a**, The intensities of Pex13-GFP (left) and mCherry-SKL (right) are cross-correlated over time to produce an image at a given lag time, ⊤ where clusters of high cross-correlation (G_XC_(0) > 0.5) indicate transient clusters of pixels where Pex13-GFP and mCherry-SKL are spatiotemporally cross-correlated. Spatial cross-correlation images at G_XC_(0) for **b**, positive control dual-labeled Pex13-GFP-mCherry expected to be highly cross-correlated across the peroxisome surface and **c**, negative control mCherry-SKL and Ant1-GFP, a peroxisome membrane protein not involved in PTS1 transport, expected to have low cross-correlation **d**, imaging Fluorescence Cross-correlation spectroscopy (iFCCS) curves taken from individual pixels demonstrate expected observations for the positive control (G_XC_(0) > 0.5) and negative control (G_XC_(0) ≅ 0). **e**, Pex13 WT strain cross-correlation at 10 minutes after Pex5 induction shows spatially (left) and temporally (right) low cross-correlation and no individual high cross-correlation pixel clusters similar to the negative control in **c**. After Pex13-GFP and SKL-mCherry associate **f**, Pex13 WT strain spatially distributed (*left*) and temporal (*right*) cross-correlation at 50 minutes after Pex5 induction shows one or more transient high G_XC_(0) > 0.5 cross-correlation clusters. **g**, Cross-sectional G_XC_(⊤) intensity (from panel **f**, *right*, dashed line) fit to a two-component Gaussian (blue, red), resolving two separate high G_XC_(0) clusters within a peroxisome. (h) Cumulative distribution of high G_XC_(0) clusters extracted from Gaussian fits (n=48) shows that 58% of pores fall within a diffraction-limited normal population while 42% are >313 nm in diameter. **i**, Image (*left*), showing at least three distinct high G_XC_(0) clusters and iFCCS temporal decays (*right*) for GFP-Pex14 and mCherry-SKL. **j**, Frequency of high G_XC_(0) clusters *versus* time following Pex5 induction shows results similar to Fig. 2 for mCherry-SKL peroxisome localization intensity in WT and S8 Pex13 mutant strains. Appearance of Pex14-mCherry-SKL high G_XC_(0) clusters following Pex5 induction is much slower, suggesting that Pex14 may form distinct condensates alone or as complexes with Pex13 with higher saturation concentrations for phase separation than those formed by Pex13. **k**, A model for peroxisome cargo transport in which Pex5-cargo protein complexes phase separate with Pex13/14 through protein-protein interactions between their IDRs to form condensates that create conduits for cargo release into the peroxisome lumen.

We first measured iFCCS of control GFP and mCherry fluorescent proteins either associated with or independent of one another to quantify the capabilities of our iFCCS analysis. We performed iFCCS with a positive control in which both GFP and mCherry are fused to

Pex13 (GFP-Pex13-mCherry, Fig. 4b). We observed a high, but incomplete, percentage of 44 ± 20% peroxisomes with ≥ 0.5 cross-correlation. Spatially, high correlation values were observed throughout the entire peroxisome, indicating that Pex13 is distributed throughout the entire membrane of the peroxisome, with correlation decays on the order of seconds (Fig. 4d, Extended Data Video 3). The total data set was comprised of > 46,000 individual correlation curves (162 peroxisomes with 289 pixels of data each), from which we calculated an average decay time based on an exponential fit to the correlation at the pixel at the center of each peroxisome of ⊤ = 2.2 ± 1.3 s, likely due to the photophysics of the fluorescent protein labels^25^. This value indicates the resolution of our measurement that we would observe in permanent- or long-lived, covalent-like association of mCherry-SKL and Pex13-GFP. A negative control of mCherry-SKL and Ant1-GFP, a peroxisomal transmembrane protein that Pex5-cargo complexes are not reported to interact with^26^, showed only 6 ± 5% peroxisomes having G_XC_(0 ms) ≥ 0.5 and the spatial images showed low correlation throughout the peroxisome with the correlation randomly scattered around G_XC_(⊤) = 0 (Fig. 4c, d, Extended Data Video 4).

We next quantified iFCCS between Pex13-GFP and mCherry-SKL at sub-peroxisome spatial scales with millisecond temporal resolutions following induction of Pex5 expression. Based on observed time-courses for appearance of mCherry foci, we collected data before and after they appeared (Fig. 2d,e). We observed that prior to mCherry-SKL import (10 minutes), the correlation is low and random throughout the peroxisome (Fig. 4e, *left*, Extended Video 5), with only 3.4 ± 0.6% peroxisomes having G_XC_(0 ms) ≥ 0.5. The temporal G_XC_ appeared similar to the negative control data (Fig. 4e, *right*). After mCherry-SKL in Pex13 WT strains (50 minutes), the number of peroxisomes with G_XC_(0 ms) ≥ 0.5 values increased over time and highly distinct clusters of pixels with G_XC_(0 ms) ≥ 0.5 were observed around the edges of the peroxisome (Fig. 4f), noticeably different than the positive control data which shows high G_XC_ throughout the entire peroxisome. These clusters of high correlation are diverse in form (ranging the entire peroxisome, polarized or centered (Extended Data Fig. 10a). Cluster sizes ranged around the diffraction limit (~300 nm), as confirmed by measuring full width half-maximal G_XC_(0) intensities across pixels centered on pixels of maximum intensity, confirming our original estimate for resolution and our supposition that this resolution would be sufficient to spatially resolve correlations of GFP and mCherry fluorescence (Fig. 4g, h). Temporal behaviors of G_XC_ were also diverse with different decays within the same peroxisome and some traces showing multiple phases (Fig. 4f right panel), along with variation decays and fluctuations observed in the decay and fluctuations of the cross-correlation over time between clusters in different peroxisomes (Extended Data Fig. 10b)., The heterogeneity over time emphasizes the transient nature of individual channels. Consequently, we did not attempt to fit the data to a model but we note that typically G_XC_ traces decayed faster than the 2.2 seconds of the positive control due to lateral diffusion of Pex13 in the membrane, suggesting that we may be capturing the faster diffusion rates of cargo through putative transient channels. Pex14-GFP showed similar G_XC_ clusters with mCherry-SKL as Pex13 but at longer times (Fig. 4i).

Finally, we observed that increases in uptake *versus* time (and thus Pex5 abundance following induction) manifested in this experiment as an increase in numbers of high G_XC_ clusters over time that reflects the increase in cargo mCherry-SKL over time observed for Pex13 and the S8 mutant (Fig. 2e,f, 4j). Surprisingly, Pex14 high G_XC_ clusters appear much later, suggesting that saturation concentrations of Pex5 and Pex14 for forming condensates must be higher (Fig. 4j).

## Discussion

The mechanism by which folded proteins and complexes are translocated into peroxisomes has been an enigma. Here we present evidence that transient biomolecular condensates composed of Pex13, Pex14, and Pex5 could act as conduits for protein import into peroxisomes and an analogous mechanism likely also occurs in chloroplasts^27^. The formation of these condensates could explain how the peroxisomal pore is formed and disassembled on demand for translocating cargos of variable size.

A biochemically-derived model for peroxin-mediated transport shows the IDR of membrane-associated Pex13 exposed to the cytosol where it captures Pex5-cargo complexes through an unknown mechanism^28,29^. It has also been proposed that Pex13 and Pex14 IDRs make long range interactions with cytosolic Pex5-cargo complexes to increase the association rate of Pex5 with the peroxisomal membrane^12^. While the Pex5 IDR has been shown to spontaneously insert into lipid membranes *in vitro*^30^ the mechanism of Pex5 interaction with the peroxisome membrane is unclear. In our model (Fig. 4k), at specific saturating concentrations, Pex13/Pex14 and Pex5-cargo complexes form a biomolecular condensate at the peroxisomal membrane which triggers LLPS in the lipid membrane and membrane insertion of Pex5. This could be followed by a reorganization of the condensate to form a transient pore in the bilayer. Such reorganization within membranes has been described before for membrane-anchored protein signaling complexes undergoing bidimensional phase separation^31^. In the lumen, the N-terminal Pex5 pentapeptide motifs could interact with the luminal Pex14-IDR domain^32^, and cargo release may be triggered by insertion of the Pex5 IDR into the Pex2-10-12 ligase channel^33^, followed by ubiquitylation of Pex5 and extraction by Pex1-Pex6 AAA ATPase to restart the cycle of cargo binding, phase separation and cargo release, as suggested recently^34^. Interestingly, blocking the Pex5 ubiquitylation step leads to accumulation of Pex13 (but not Pex14)^33^, suggesting that Pex5 recycling may be linked to resetting Pex13 homeostasis on the peroxisomal membrane. Proteomics studies detect Pex13 ubiquitylation (K233, K242)^35^ and phosphorylation (S246, S251)^36^ at sites predicted to interact with Pex14 (residues 236-246 of Pex13). It remains to be determined whether posttranslational modifications regulate phase separation of Pex13 and thereby membrane interactions with Pex14-Pex5.

Our model serves as a framework for advancing knowledge on the function and pathology of peroxisomes. A number of questions about protein transport can now be addressed in the context of our model. For instance, it will be of interest to determine if, like NPCs, Pex13/14 condensates act as selective filters, excluding proteins above a critical size that do not interact with Pex5. The existence of Pex5 and Pex14 channels at different concentrations of Pex5 suggests an analogy to active transporters of transition metals where high and low affinity metal transporters function to transport when extracellular metal concentrations are low or high, respectively^37–39^. Our model is consistent with PTS1 import required for normal peroxisome biogenesis. For instance, import of Pex8p requires only Pex5 and Pex14^40^, the transport of PTS1- and PTS2-proteins occurs through distinct aqueous protein-conducting channels^3,41^, and over-expression of Pex5p can partially compensate an import defect of a PEX14 null mutant, supporting the notion that increasing in saturating concentrations for phase separation of channel-forming proteins in mutant peroxins could be the cause of protein transport deficits^42^

Defects in the import machinery of the peroxisome in humans results in peroxisome biogenesis disorders, the most severe being Zellweger spectrum disorders (ZSD). Mutations in Pex5, Pex13 and Pex14 that lead to their deficiency or total absence have been identified in ZSD patients^43,44^. Interestingly, although the IDRs have many sequence variations, none of these contribute to disease. Perhaps this is because phase separation remains robust to such sequence variations. Finally, given that Pex13, Pex14 and Pex5 are broadly conserved, the mechanism of transport we propose is likely conserved across eukaryotes^45^. Notably, the patterning of aromatic residues in Pex13 is conserved from yeast to human, despite very low levels of sequence conservation (Extended Fig.2b).

Future work could pursue resolving the quantitative dynamics of the transient pores with increased resolution beyond our iFCCS capabilities. Traditional confocal FCCS or wide field imaging FCCS with higher frame rates using recent innovations in scientific CMOS technologies could expand the range to μs-minute dynamics.

## Methods

### Strains, gene editing and culture conditions

The yeast strains used in this study are listed in Table S1. BY4742 *MATα his3Δ1 leu2Δ0 lys2Δ0 ura3Δ0* background or seamless SWAT background (*MATα his3Δ1 leu2Δ0 met15Δ0* ura3Δ0 lys+ *can1Δ*∷*pGAL1-SceI*∷*pSTE2-SpHIS5 lyp1Δ*∷*pSTE3-LEU2*) in the case of Pex13-GFP and GFP-Pex14 strains^1^. Yeast transformation, manipulation of *Escherichia coli*, and the preparation of bacterial growth medium were performed as described previously^2^. Gene editing was performed by CRISPR-Cas9 editing according to Shaw *et al*.^3^. Imaging experiments were performed with exponentially growing cells in synthetic complete media. A vector constitutively expressing mCherry-SKL was designed by PCR amplification of a p413 ADH1 mCherry template and subcloned into p413 ADH1 at *Xba*I and *Xho*I restriction sites. For kinetics and single-molecule microscopy studies of mCherry-SKL import, The *GAL1*promoter (p*GAL1*) was amplified from the genome of BY4742 strain and inserted upstream of the *PEX5* gene by CRISPR/Cas9 editing. Plasmids used in this study are listed in Table S2.

### Induction of Pex5 expression and analysis

Pre-cultures of strains expressing pGal1-Pex5 were grown in synthetic complete medium lacking histidine (SC-His) containing 2% glucose w/v overnight at 30°C with shaking. The preculture was washed, diluted to O.D._600_ of 0.2 and incubated overnight in SC-His medium containing 2% raffinose. The overnight culture was diluted again in SC-His medium containing 2% raffinose and allowed to reach an O.D. _600_ of 0.6. Cells were incubated on glass bottom 8-well microscopy chambers (Lab-Tek Nunc) coated with 0.1% concanavalin A w/v (Sigma-Aldrich ConA # C-7275) for 20 minutes. Unbound cells were washed once with SC-His raffinose medium followed by addition of SC-His medium containing 2% galactose w/v to induce Pex5 expression. Images were acquired with a Quorum Diskovery platform consisting of a spinning disc confocal system (see microscopy) using a 100X oil immersion objective.

### Protein purification

Bacterial codon-optimized sequences were generated for all Pex13, Pex14 and Pex5 constructs by Bio Basic Life Science. To generate fusion proteins with glutathione-S-transferase (GST), Pex14-IDR, Pex5 and Pex13-IDR WT and mutants oligonucleotide were subcloned into an in-house pGEX-TEV vector between BamHI and EcoRI sites. An N-terminal cysteine residue was added to the Pex13 and Pex14 IDRs, to allow fluorescent labeling via maleimide chemistry. The mCherry-SKL-coding oligonucleotide was subcloned into pET15b to generate a 6xHis-tagged fusion protein. Proteins were expressed in *E. coli* BL21 strain (Novagen) in LB-Amp medium. Protein expression was induced at OD_600_ = 0.8 with 0.6 mM IPTG at 30°C for 4 h. Cells were collected by centrifugation, resuspended in lysis buffer (20 mM Tris pH 7.4, 0.2 mM EDTA, 1 M NaCl, 1 mM DTT) and lysed by passing the suspension through a French press twice, alternating with sonicating and homogenizing cycles. Cell lysates were centrifuged at 105 000 × g for 1 h at 4°C, and the resulting supernatant was incubated with glutathione (GSH) sepharose resin for 1 h at room temperature with gentle shaking. Supernatant was filtered to collect resin-bound GST-fusion protein and washed 4X with lysis buffer, 2X with TEV buffer (25 mM Tris pH 7.4, 100 mM NaCl, 5 mM DTT). The GST tag was cleaved by incubating the resin in TEV buffer containing TEV enzyme. The sample was centrifuged after incubation with TEV and the supernatant containing the cleaved protein was purified by anion exchange chromatography using a column packed with either Q Sepharose High Performance (GE Healthcare) for Pex13 and 14 IDRs or SP Sepharose High Performance (GE Healthcare) for Pex5. The column (50 mL) was equilibrated with buffer A (20 mM Tris, pH 7.4, 2.5 mM DTT) and the protein was eluted using a gradient of Buffer B (0 to 1 M NaCl over 0.5 L). Fractions containing the protein were pooled and further purified with gel filtration using a Superose-12 column equilibrated with 20 mM Tris-HCl pH 7.4, 500 mM NaCl. IDRs were eluted under denaturing conditions (4 M urea added to buffer A) and buffer exchanged in 4 M guanidinium hydrochloride before flash freezing in liquid nitrogen and storing in aliquots at −80°C. To purify mCherry-SKL, cell lysates were prepared using the above mentioned method and buffers with the following changes. Supernatant of cell lysate was incubated with Chelating Sepharose™ fastflow sepharose resin (GE Healthcare) charged with Ni^2+^ (according to the manufacturer’s protocol) for 1 h at room temperature with gentle shaking. Resin-bound 6XHis-fusion protein was washed 4X with wash buffer (20 mM Tris, pH 7.4, 500 mM NaCl, 5 mM Imidazole) and eluted using an imidazole gradient. The 6XHis tag was cleaved by incubating the resin in wash buffer containing Thrombin enzyme. The supernatant containing untagged mCherry-SKL was purified by anion exchange chromatography and gel filtration as mentioned above.

### *In vitro* phase separation assays

Screening to determine the saturation concentration of protein was done in 384 well microplates (Corning). A small volume (0.5 μL) of concentrated protein fractions in 4 M Guanidine hydrochloride was placed on the imaging chamber and diluted rapidly in phase separation buffer (20 mM Tris pH 7.4 containing 10% PEG 8000). Images were taken immediately post dilution for testing droplet formation. Pex13 fusion events were captured by 30 ms time lapses using a 60X objective on a Nikon Eclipse TE2000E microscope. Fusion of two droplets of similar size was analyzed as previously described^4^ with required changes to the MATLAB code for analyzing brightfield images. For reconstitution assays, Pex5 was labeled with AF dye 647 (Click Chemistry Tools) according to manufacturer’s protocol. A small amount of labeled Pex5 protein was added to unlabeled protein (2% labeled protein) and tested for droplet formation to determine whether labeling affects saturation concentration for phase separation. Equimolar concentration of Pex5-AF-647 and mCherry-SKL was mixed in a microfuge tube taking care to avoid introduction of bubbles and incubated at room temperature for 30 minutes. Pex13-IDR stock (3 mM) was rapidly diluted to 10 μM in phase separation buffer directly on glass bottom microslides (Ibidi) and formation of condensates was tested. Pex5-AF-647 and mCherry-SKL complex was added to the preformed Pex13 droplets to reach a final concentration at which Pex5 and mCherry-SKL do not form condensates (1 μM) (Extended Fig 8c). Control reactions lacking either mCherry-SKL or Pex5 were prepared and analyzed simultaneously. Condensates were imaged immediately with the Quorum Diskovery platform consisting of a spinning disc confocal system (see Microscopy and analysis) using a 100X oil immersion objective.

### Microscopy and analysis

In vivo imaging experiments and imaging of fluorescently labelled protein droplets were performed using the Quorum Diskovery platform consisting of a Leica DMi6000 inverted microscope equipped with a Spectral Diskovery multimodal imaging system attached to either a HamamatsuEM×2 camera or ORCA FLASH 4.0 V2 digital complementary metal-oxide-semiconductor camera. Metamorph software (Molecular Devices) was used for image acquisition controls. Fluorescence images were acquired in a series of *z*-axis planes (0.5 μm apart). Fluorescence intensity measurement was performed using *z*-axis projection of images and the maximum fluorescence in nine adjacent pixels was measured using Fiji software^5^. Screening and brightfield imaging of protein droplets was done using a Nikon Eclipse TE2000E microscope equipped with a 60X/1.45 plan objective (Nikon) and a Cool SNAP HQ camera.

### Fluorescence Recovery After Photobleaching (FRAP)

FRAP was performed to determine recovery of mCherry-SKL partitioned within Pex13-IDR droplets in the presence of Pex5. Images were acquired with spinning disc confocal system (Quorum) using a 100X oil immersion objective and the FRAP module of Metamorph software. Five control images were taken using the mCherry filter prior to bleaching. A region of interest (ROI) of 1 μm diameter was set for photobleaching with a 488 nm laser at 100% power for 150 ms. Images were acquired immediately every 10 s for 1 min, then every 15 s for another minute and every 30 s for the following 3 minutes (total 300 seconds). Relative fluorescence intensity of the ROI (F) through all the stacks was measured using Fiji image processing package^5^. Additionally, recovery in a nearby unbleached fluorescent droplet (control), and a non-fluorescent background region (F_b_) was also measured. The photobleaching rate (r) was calculated by comparing the fluorescence of the control region before (F_c0_) and after (F_c_) photobleaching. r = F_c_/F_c0_. The fluorescence intensity of droplet ROI was normalized with the background and photobleaching rate as follows: F_norm_ = (F - F_b_) / r. Finally, the normalized fluorescence intensity was curve fit with a one-phase exponential equation of Graphpad Prism software.

### Imaging Fluorescence Cross-Correlation Spectroscopy (iFCCS)

The output of the microscopy data is two image stacks corresponding to 1200 pixels × 1200 pixels (84 μm × 84 μm) and 150 frames (22.5 s) in size (Extended Data Fig. 9). A corresponding image of fiduciary markers (Thermo Fisher T7279) taken at the coverslip/sample interface of the same area of the sample is also collected. The analysis first uses the fiducial markers to align the green and red channels using MATLAB’s built in control point selection and correlation functions to ensure pixel-level accuracy between channels. The centroid position of at least six fiducial markers are selected by hand (Extended Data Fig. 9b) and the coordinates exported (MATLAB’s ‘cpselect’). The points are further refined by using a normalized cross correlation between the red and green channels (‘cpcorr’). The corresponding points are used to calculate an affine transformation to shift the green channel of the Pex13-GFP image stack with respect to the mCherry-SKL data. Next, the centroid locations of peroxisomes of interest are localized using a previously established particle localization analysis (Extended Data Fig. 9c, dashed circle provided as guide for the eye)^6^. The location of peroxisomes is identified from the sum of all collected frames. Single particle tracking analysis has shown that diffusion of the peroxisomes over one pixel during observation time (>2.2 × 10^4^ μm^2^/s) in negligible. The GFP and mCherry intensities of the surrounding 17 pixels × 17 pixels of the peroxisome centroid location are extracted (Extended Data Fig. 9d, intensities of centroid pixel boxed in 9c). The intensity fluctuations, δF(t), relative to the mean intensity are calculated according to δF(t) = F(t)-<F(t)> and used to calculate the cross correlation, G(⊤), via MATLAB’s ‘xcorr’ function with a normalization based on the autocorrelations of the GFP and mCherry signals at zero lag will be equal to 1 (MATLAB’s ‘coeff’). The resulting cross-correlation data can be used to determine protein-protein interactions at a specific locations and time. Viewed spatially, the magnitude of the cross-correlation at ⊤=0 and subsequent lags can localize where Pex-13 and mCherry-SKL are at relative to each other at sub-organelle resolutions (Extended Data Fig. 9f; dashed line indicates same area as shown in Fig 9c). Temporally (Extended Data Fig. 9g), the cross-correlation decay curves determine how the mCherry fluorescence changes with respect to the GFP fluorescence at an individual pixel. The magnitude, shape, and decay time of the curve can be used to quantify dynamics.

## End Notes

## Acknowledgements

The authors thank Maya Schuldiner for seamless SWAT strains and advice on the *VPS1* knockouts, Jackie Vogel for providing strains from the YKO collection, Edward Lemke for suggestions regarding in vitro phase separation experiments, Haytham Wahba and James Omichinski for sharing expertise in protein purification, Tina Tai for making the mCherry-SKL construct, Philippe Garneau for technical assistance and Nicolas Stifani for help with FRAP. The authors acknowledge support from Canadian Institutes of Health Research (CIHR) grant MOP-GMX-152556 and Human Frontier Science Program grant RGP0034/2017 (SWM) and cross-disciplinary fellowship (LT000805/2018-C) (IOLB), the Case Western Reserve College of Arts and Sciences and the Department of Physics (LK and ZZ) and NIH NIGMS grant R35GM12466 (LK) for financial support of this work.

## Authors contributions

SWM and IOLB conceptualized this study. IOLB generated and characterized yeast strains for induction of Pex5 expression, standardized microscopy conditions for iFCCS and Pex5 induction and designed constructs for protein purification. RR performed Pex5-cargo transport experiments, purified proteins, standardized and performed in vitro experiments and acquired data for iFCCS. XCG analyzed Pex13 droplet fusion data. ZZ and LK generated pipelines for iFCCS and compiled the resulting data. RR, IOLB, XCG, LK and SWM analyzed and interpreted data. RR, IOLB, LK and SWM wrote the manuscript and all authors provided feedback and editorial support.

## Data availability statement

All data generated or analyzed during this study are included in this published article (and its supplementary information files)

## Extended Data

**Extended Data Fig. 1.**
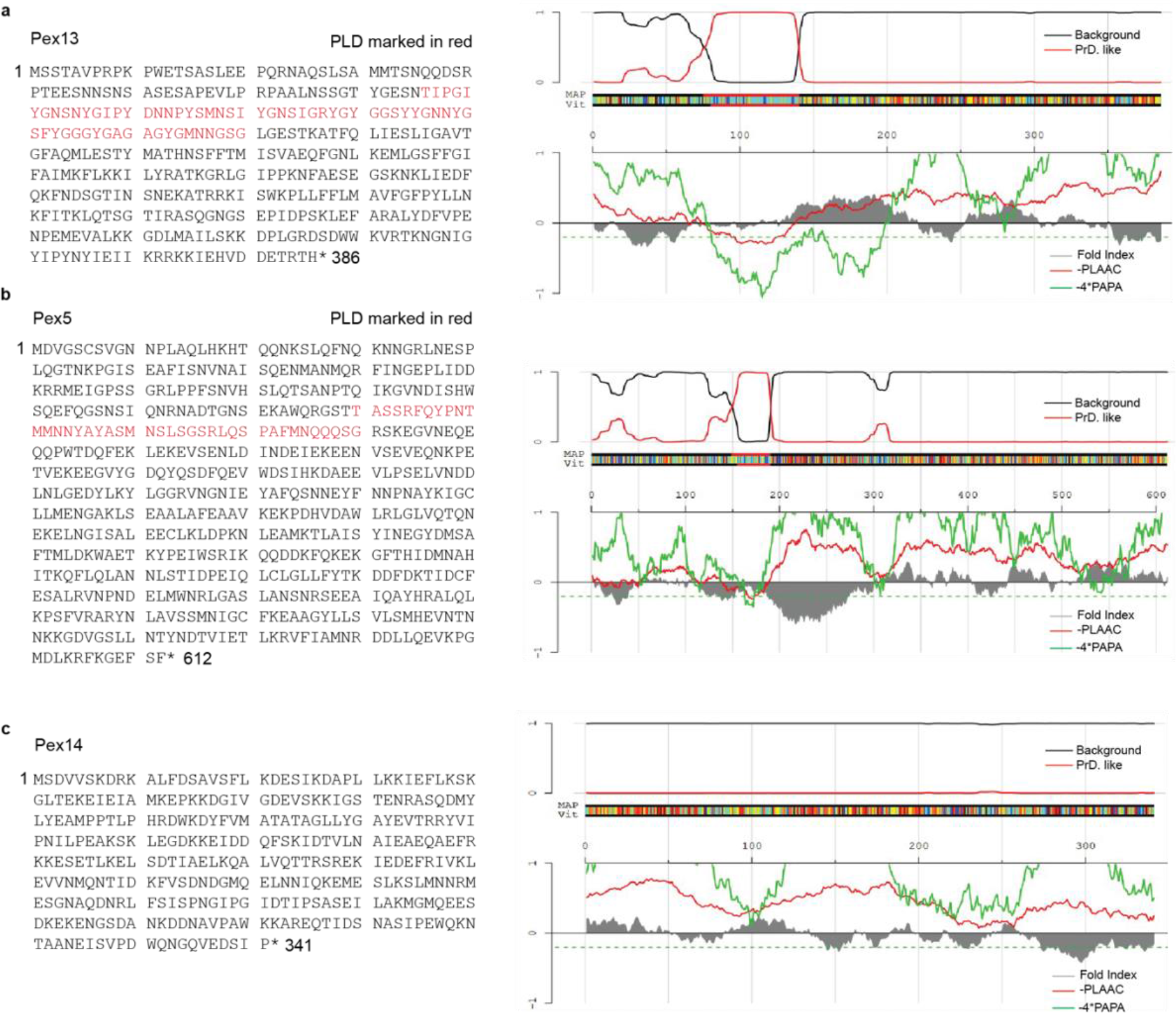
Pex13 and Pex5 have prion-like domains (PLD). Analysis of **a**, Pex13 **b**, Pex5 and **c**, Pex14 sequences for the presence of PLD using PLAAC^1^. Predicted PLD sequence is marked in red for Pex13 and Pex5 on the left, corresponding to the outputs on the right. Pex14 does not have a predicted PLD.

**Extended Fig. 2.**
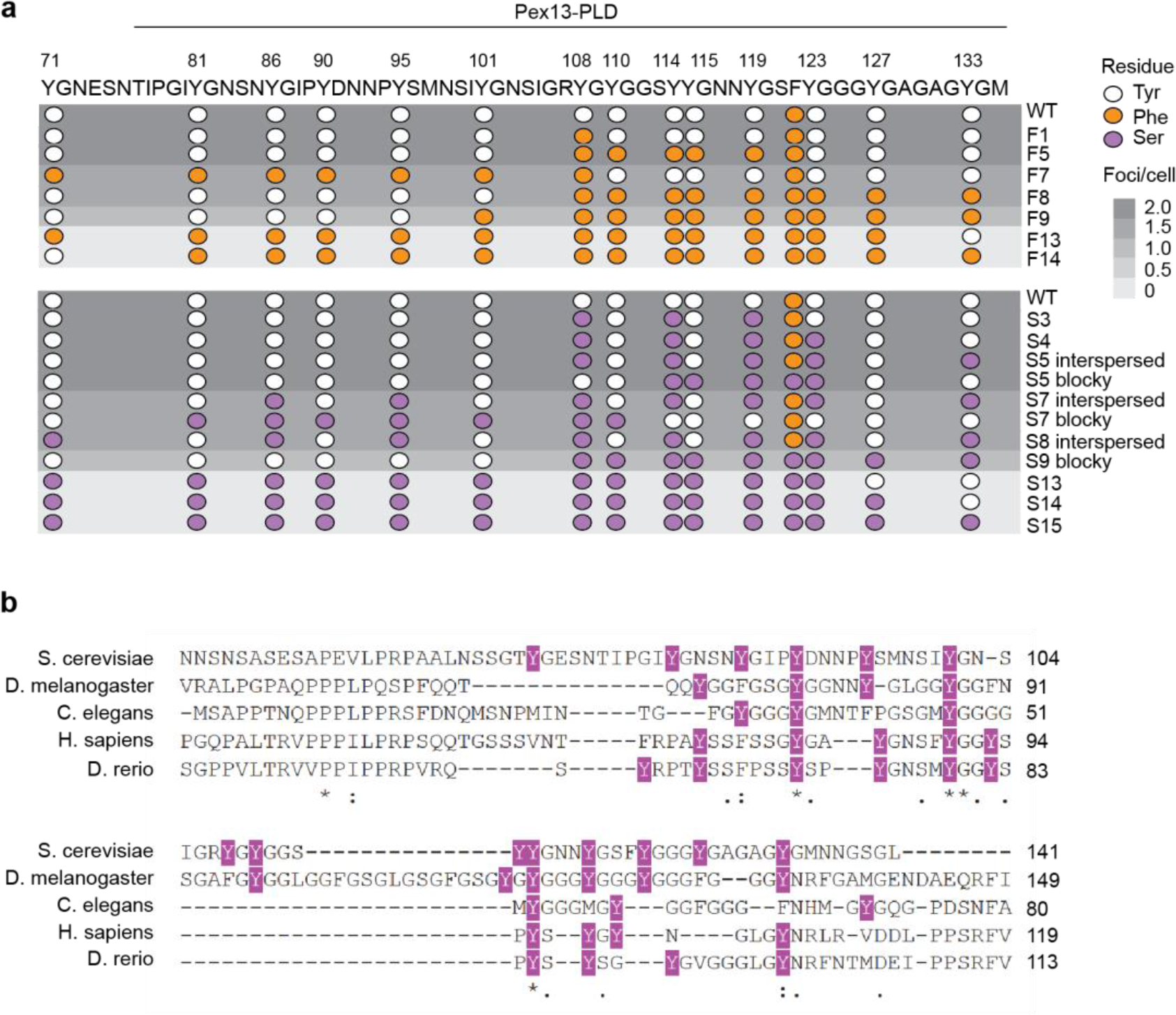
Pex13 Tyr to Phe or Ser mutants generated by CRISPR-Cas9 gene editing. **a**, Distribution of Tyrosine (Y, white) to Phenylalanine (F, orange) or Serine (S, purple) point mutations introduced in the Pex13 PLD using CRISPR-Cas9 editing. Note that the S5 blocky, S9, S13, S14 and S15 mutants additionally include the replacement of an F at residue 122 to S. Point mutation heat map shown in shades of gray denotes the number of mCherry foci observed in the indicated cells. n > 100 cells were sampled for data pooled from three independent experiments. **b**, Multiple sequence alignment of the Pex13 sequence from the indicated five species, showing conserved Tyr residues within the predicted yeast PLD sequence.

**Extended Data Fig. 3.**
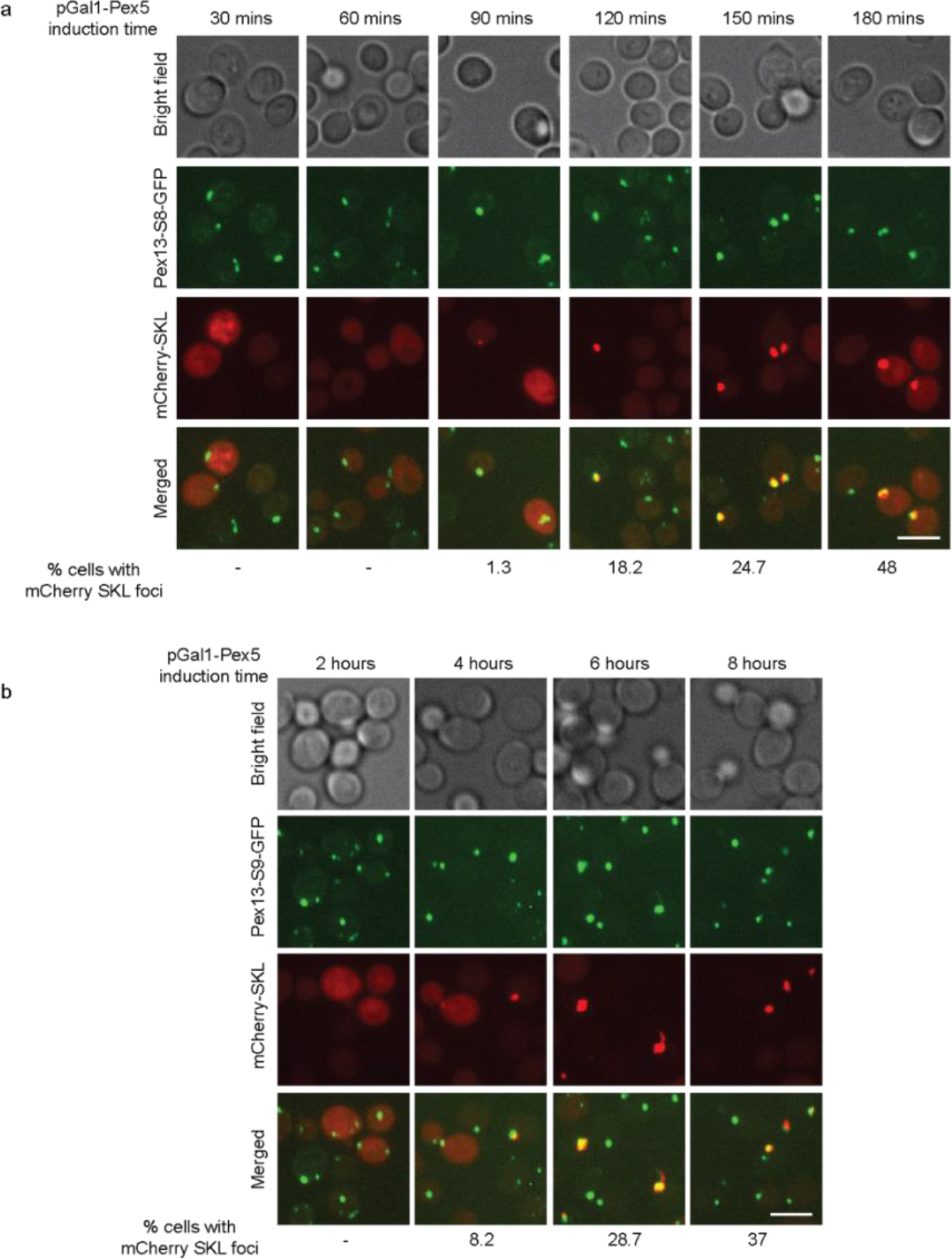
mCherry-SKL transport kinetics in Pex13-S8 and Pex13-S9 mutants. Spinning disc confocal images of **a**, Pex13-S8-GFP and (b) Pex13-S9-GFP cells taken at the indicated time points after induction of Pex5. Scale bar 5 μm. Percentage cells displaying mCherry-SKL foci are indicated below each panel. N = 50 cells per sample.

**Extended Data Fig. 4.**
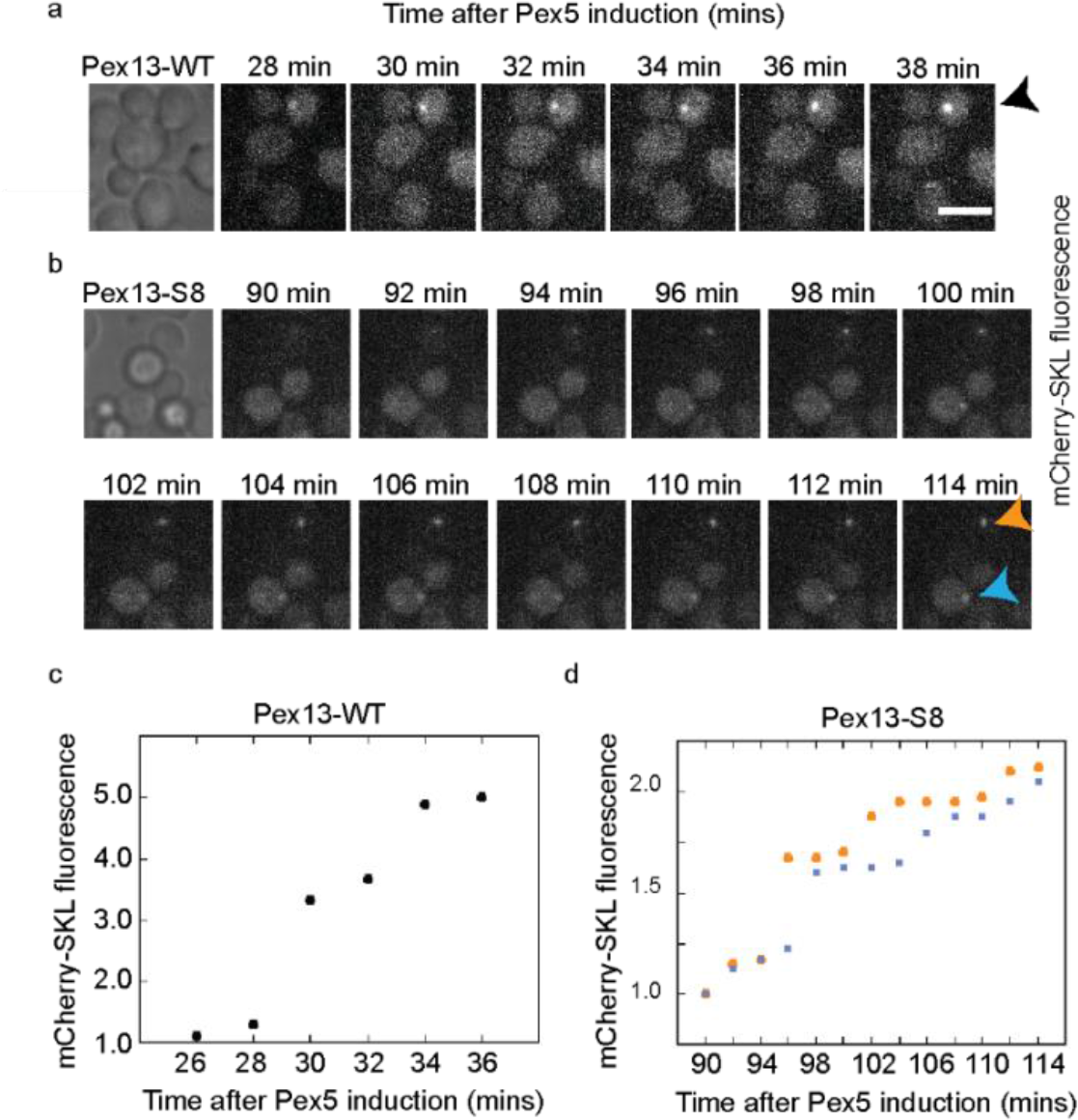
mCherry-SKL cargo transport kinetics in Pex13 WT and Pex13 Tyr to Ser mutants. Timelapse imaging to follow formation of mCherry-SKL foci in **a**, Pex13 WT cells (black arrow) and **b**, Pex13 S8 mutant cells (orange and blue arrows). Spinning disc confocal images were taken every 2 mins after induction of Pex5 expression at the indicated timepoints. Scale bar 5 μm. **c,d**, mCherry-SKL foci fluorescence intensity of the cells imaged in panels **a** and **b**, respectively, plotted as a function of time after induction of Pex5 expression.

**Extended Data Fig. 5.**
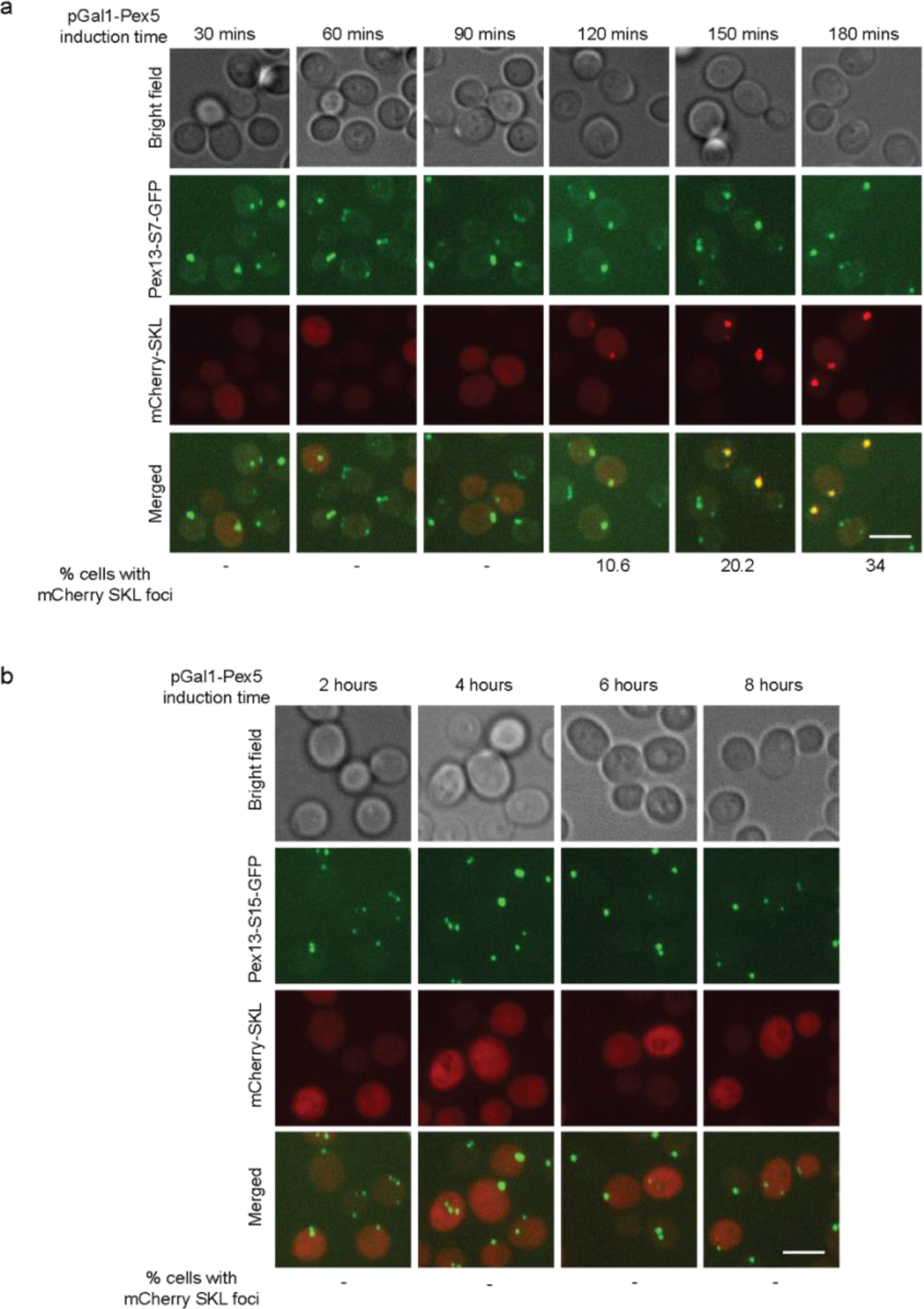
mCherry-SKL transport kinetics in Pex13 S7 and Pex13 S15 mutants. Spinning disc confocal images of **a**, Pex13-S7-GFP and **b**, Pex13-S15-GFP cells taken at the indicated time points after induction of Pex5. Scale bar 5 μm. Percentage cells displaying mCherry-SKL foci are indicated below panel a. mCherry-SKL remains cytoplasmic in Pex13 S15 cells throughout the duration of the experiment (8 hours). n = 50 cells.

**Extended Data Fig. 6.**
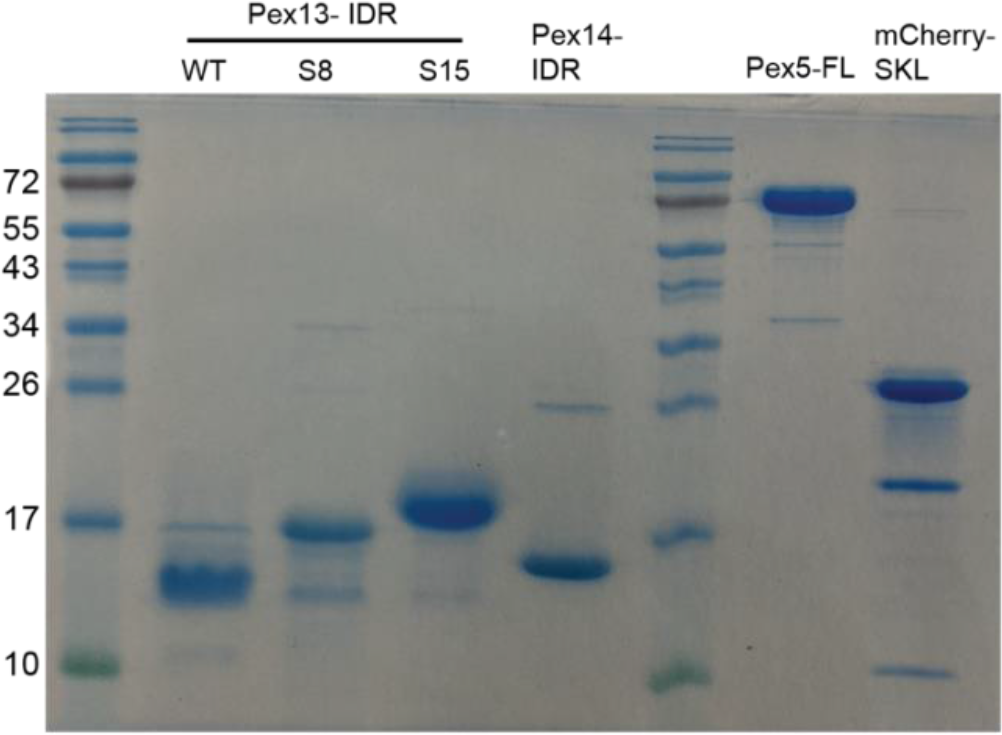
Pex protein purification. 1 μg of the indicated purified proteins were loaded into wells of 15% SDS-PAGE gels and electrophoresis was carried out in Tris glycine buffer at 150 volts for 1.5 hours, followed by staining with Coomassie stain.

**Extended Data Fig. 7.**
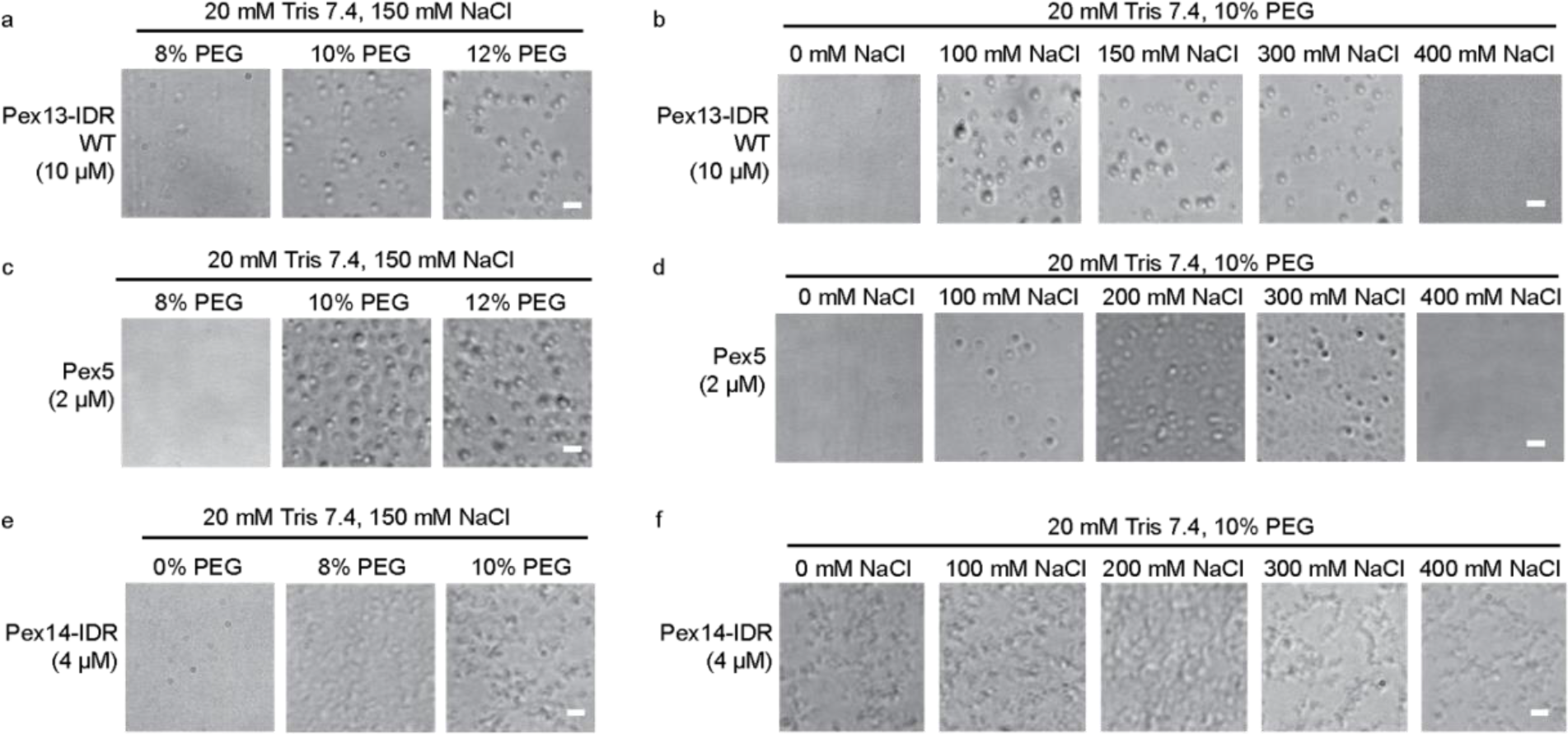
*In vitro* phase separation of PTS1 peroxins. Brightfield images of condensates formed by **a,b**, Pex13-IDR WT **c,d**, full length Pex5 and **e,f**, Pex14 IDR. Phase separation Pex13-IDR WT and full-length Pex5 are dependent on PEG and salt concentration. Pex14-IDR forms elongated particles dependent on PEG but independent of salt in the phase separation buffer (20 mM Tris pH 7.4 containing 10% PEG 8000). Scale bar 5 μm.

**Extended Data Fig. 8.**
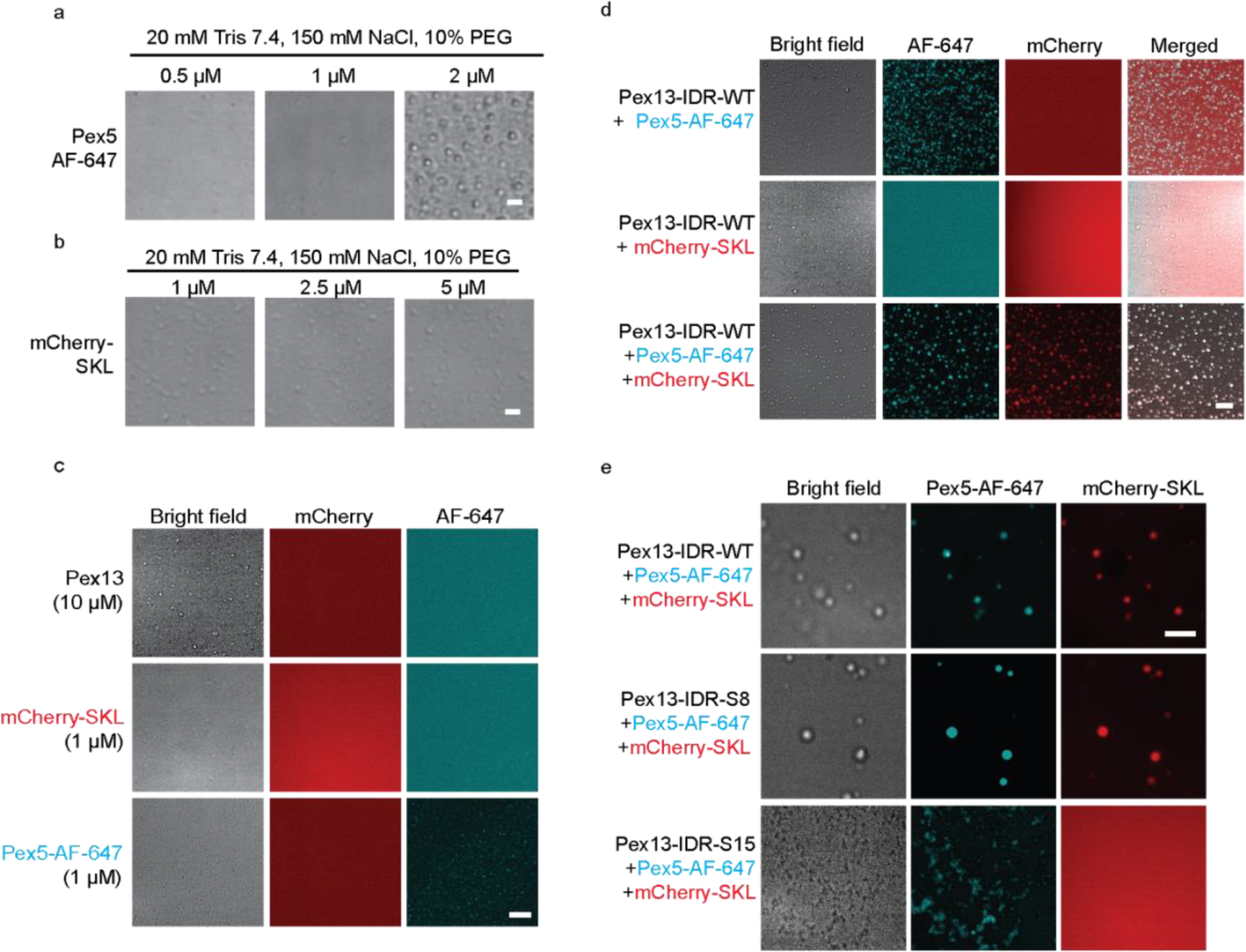
mCherry-SKL partitioning into Pex13 condensates is Pex5-dependent. Titration of **a**, Pex5 and **b**, mCherry-SKL to determine the concentration at which they do not form condensates. 1 μM of both Pex5 and mCherry-SKL were used for *in vitro* reconstitution assays with Pex13-IDR. **d**, Controls showing that under the conditions used for reconstitution assays, only Pex13 formed visible condensates. **d**, Reconstitution experiments to measure partitioning of Pex5 labelled with AF 647 dye and mCherry-SKL into Pex13-IDR WT condensates immediately after mixing at room temperature. **e**, Pex5 and mCherry-SKL partitions into Pex13-IDR S8 and Pex13-IDR S15 does not form condensates. Scale bar 5 μm.

**Extended Data Fig. 9.**
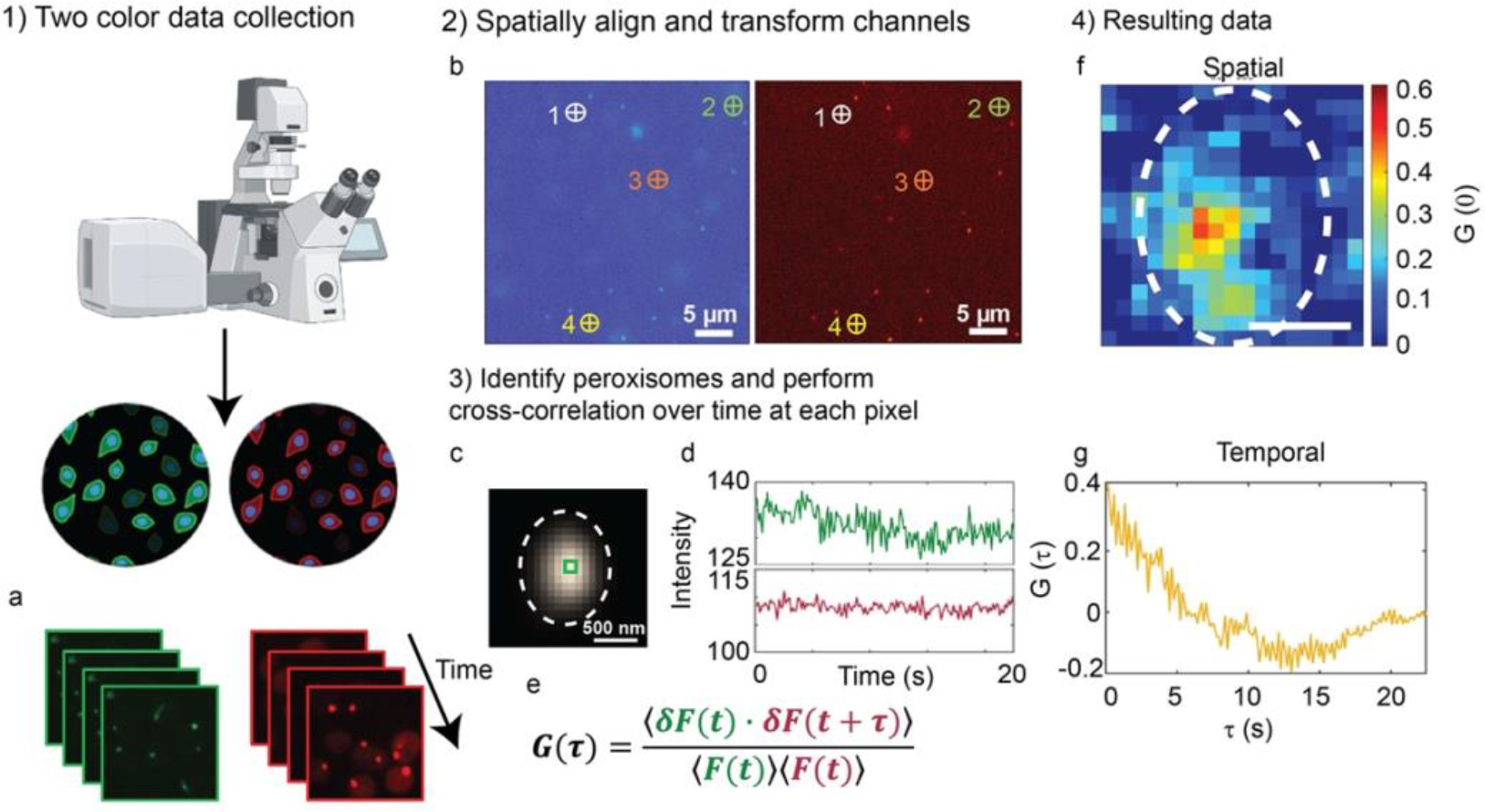
Imaging fluorescence cross-correlation spectroscopy (iFCCS) determines where mCherry-SKL and GFP-Pex13 intensities correlate within individual peroxisomes and how the relative intensities change over milliseconds-seconds time scales. **a**, The output of the microscopy data is two image stacks corresponding to 1200 pixels × 1200 pixels (84 μm × 84 μm) and 150 frames (22.5 s) in size. A corresponding image of fiduciary markers taken at the coverslip/sample interface of the same area of the sample is also collected. The analysis first uses the fiducial markers to align the green and red channels using MATLAB’s built in control point selection and correlation functions to ensure pixel-level accuracy between channels. **b**, The centroid position of at least six fiducial markers are selected by hand and the coordinates exported (MATLAB’s ‘cpselect’). The points are further refined by using a normalized cross correlation between the red and green channels (‘cpcorr’). The corresponding points are used to calculate an affine transformation to shift the green channel of the Pex13-GFP image stack with respect to the mCherry-SKL data. **c**, Next, the centroid locations of peroxisomes of interest are localized using a previously established particle localization analysis (dashed circle provided as guide for the eye)^2^. The location of peroxisomes are identified from the sum of all collected frames. Single particle tracking analysis has shown that diffusion of the peroxisomes over one pixel during the observation time (>2.2 × 10^4^ μm^2^/s) in negligible. **d**, The GFP and mCherry intensities of each of the surrounding 17 pixels × 17 pixels of the peroxisome centroid location are extracted. The intensities of centroid pixel boxed in **c** are shown in **d**. Using the equation in **e**, the intensity fluctuations, δF(t), relative to the mean intensity are calculated according to δF(t) = F(t)-<F(t)> and used to calculated the cross correlation, G(⊤), *via* MATLAB’s ‘xcorr’ function with a normalization based on the autocorrelations of the GFP and mCherry signals at zero lag will be equal to 1 (MATLAB’s ‘coeff’). The resulting cross-correlation data can be used to determine protein-protein interactions at a specific locations and time. **f**, Viewed spatially, the magnitude of the cross-correlation at ⊤=0 and subsequent lags can localize where Pex-13 and mCherry-SKL are at relative to each other at sub-organelle resolutions; dashed line indicates same area as shown in (c). **g**, Temporally, the cross-correlation decay curves determine how the mCherry fluorescence changes with respect to the GFP fluorescence at an individual pixel. The magnitude, shape, and decay time of the curve can be used to quantify temporal dynamics.

**Extended Data Fig. 10.**
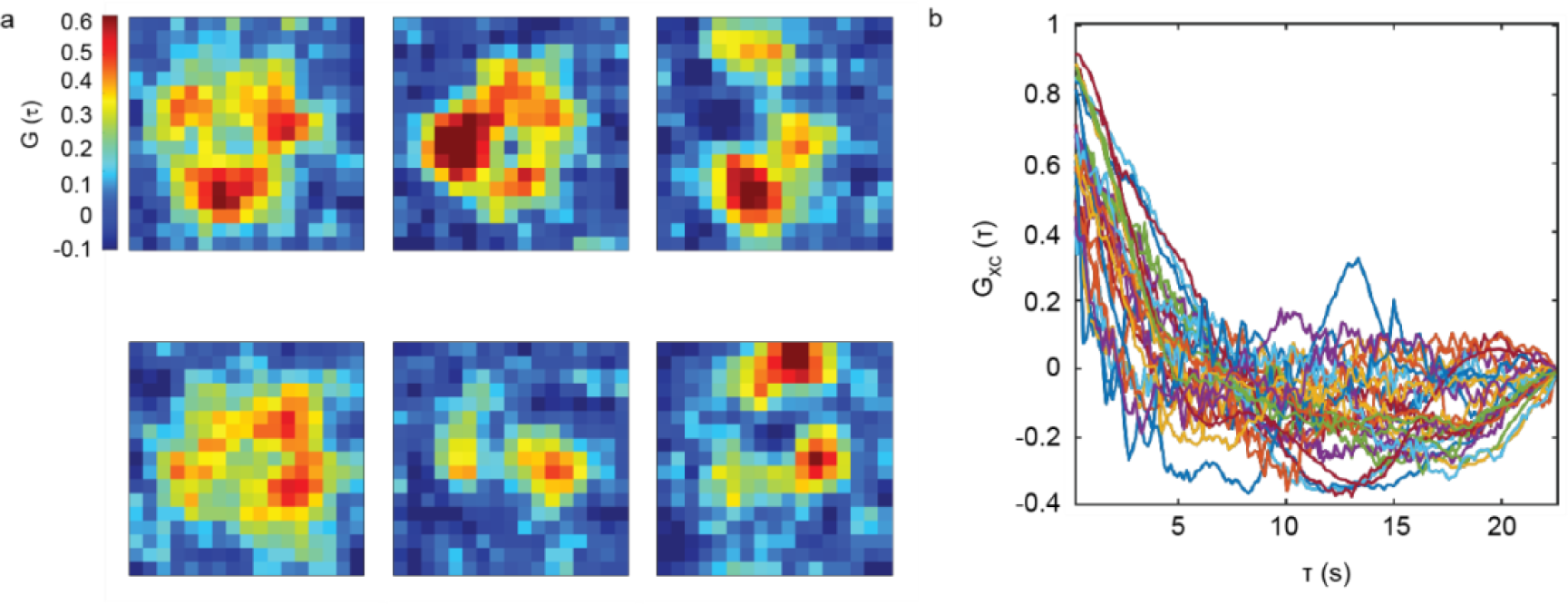
iFCCS spatial and temporal data shows transient clusters of correlated signal of mCherry-SKL and GFP-Pex13. **a**, Spatial iFCCS data from six different peroxisomes in WT 50 min after Pex5 induction where distinct transient clusters of correlated signal are observed. Images measure 1.19 μm × 1.19 μm in size. **b**, Transient nature of individual clusters are observed in the decay and fluctuations of the cross-correlation over time between clusters in different peroxisomes (n=27 peroxisomes).

**Extended Data Video 1 | Pex13-IDR WT condensate fusion**. Timelapse of Pex13-IDR WT condensates imaged at 30 ms intervals. Scale bar 5 μm. Frame rate 5 frames per second.

**Extended Data Video 2 | Pex13-IDR S8 condensate fusion**. Timelapse of Pex13-IDR S8 condensates imaged at 30 ms intervals. Scale bar 5 μm. Frame rate 5 frames per second.

**Extended Data Video 3 | Cross-correlation video of positive control shows high spatial correlation between dual-labeled Pex13-GFP-mCherry**. A peroxisome identified by single particle tracking algorithm is located at the center of the image which measures 1.19 μm × 1.19 μm in size. The time listed above the figure indicates the lag time and the color indicates the magnitude of the cross-correlation at the given lag time. Note the extended G_XC_(⊤) scale from 0 to 1 compared to the scales used in Extended Videos 4, 5, and 6.

**Extended Data Video 4 | Cross-correlation video of negative control shows low spatial correlation between mCherry-SKL and Ant1-GFP**. A peroxisome identified by single particle tracking algorithm is located at the center of the image which measures 1.19 μm × 1.19 μm in size. The time listed above the figure indicates the lag time and the color indicates the magnitude of the cross-correlation at the given lag time. Data obtained from a peroxisome identified 50 min after induction, the same time as the wild type data shown in Extended Video 6.

**Extended Data Video 5 | Cross-correlation video of Pex13 WT cells 10 min after induction shows low spatial correlation between SKL-mCherry and Pex13-GFP**. Data is similar to that of the negative control showing low correlation. A peroxisome identified by single particle tracking algorithm is located at the center of the image which measures 1.19 μm × 1.19 μm in size. The time listed above the figure indicates the lag time and the color indicates the magnitude of the cross-correlation at the given lag time.

**Extended Data Video 6 | Cross-correlation video of wild type 50 min after induction shows higher spatial correlation between SKL-mCherry and Pex13-GFP distributed around the edge of peroxisome**. Three distinct pores can be visualized in the first frame of the video which then quickly decays as lag time increases indicating the transient nature of the pores. A peroxisome identified by single particle tracking algorithm is located at the center of the image which measures 1.19 μm × 1.19 μm in size. The time listed above the figure indicates the lag time and the color indicates the magnitude of the cross-correlation at the given lag time.

**Table S1.**
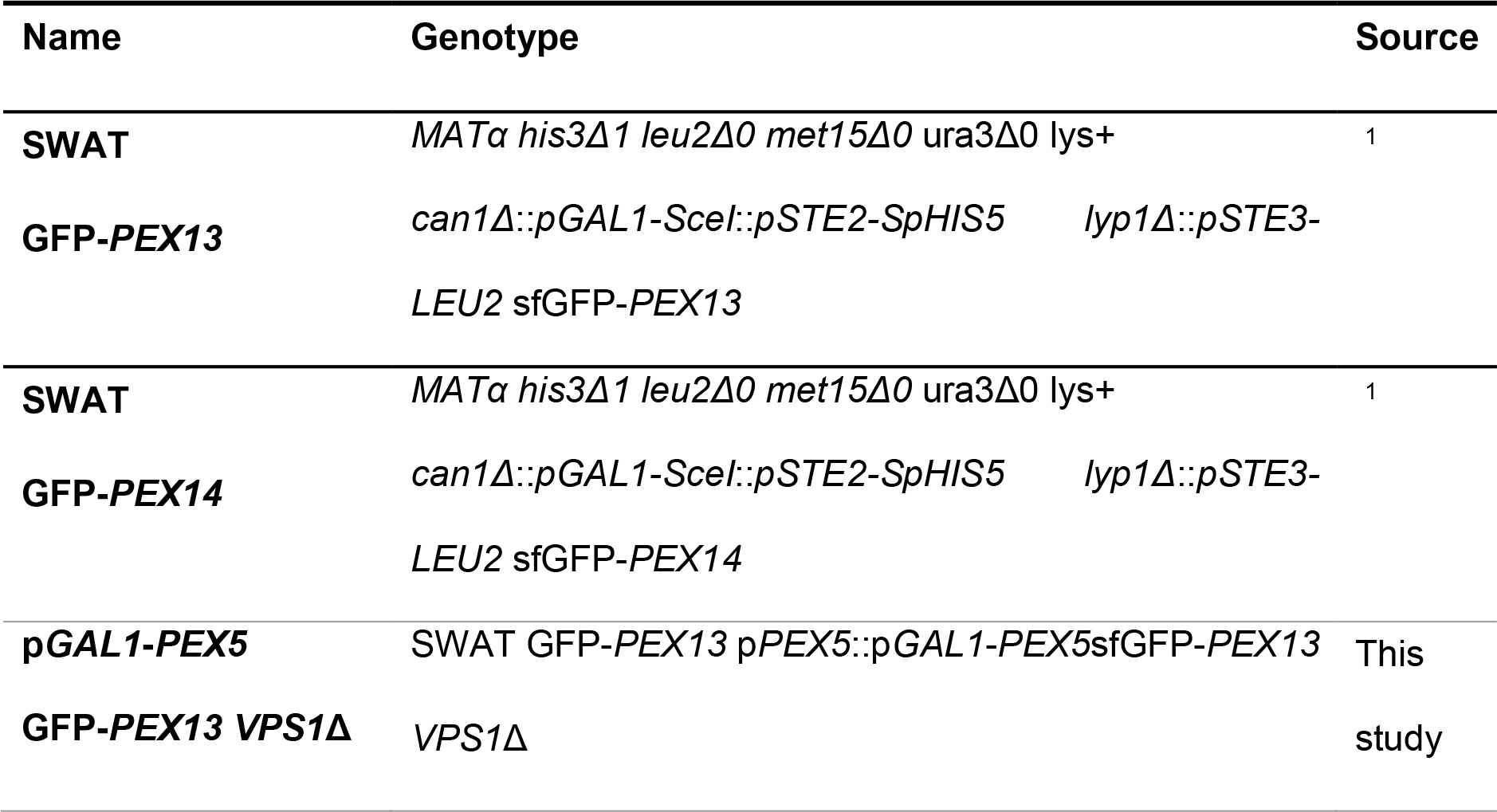
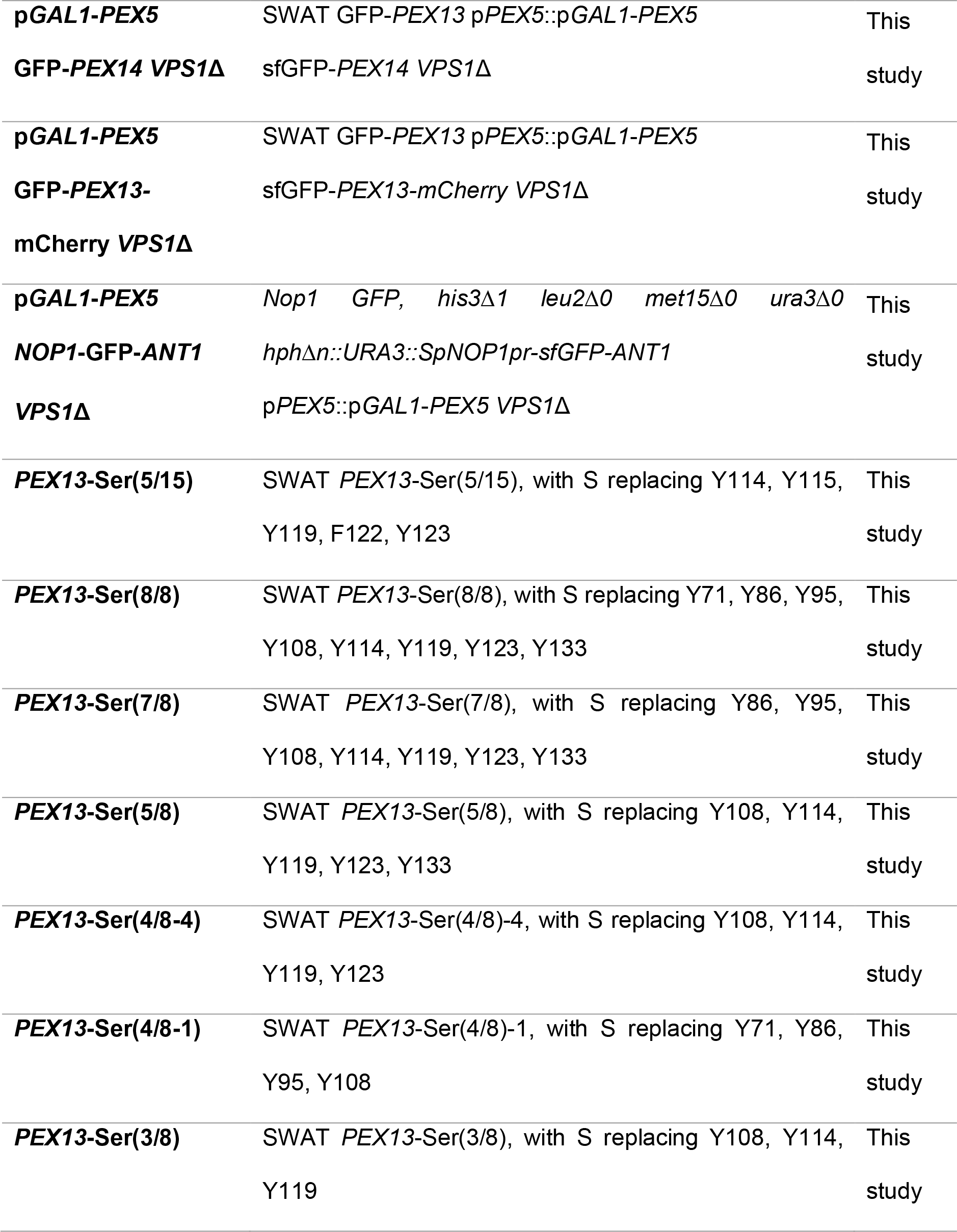
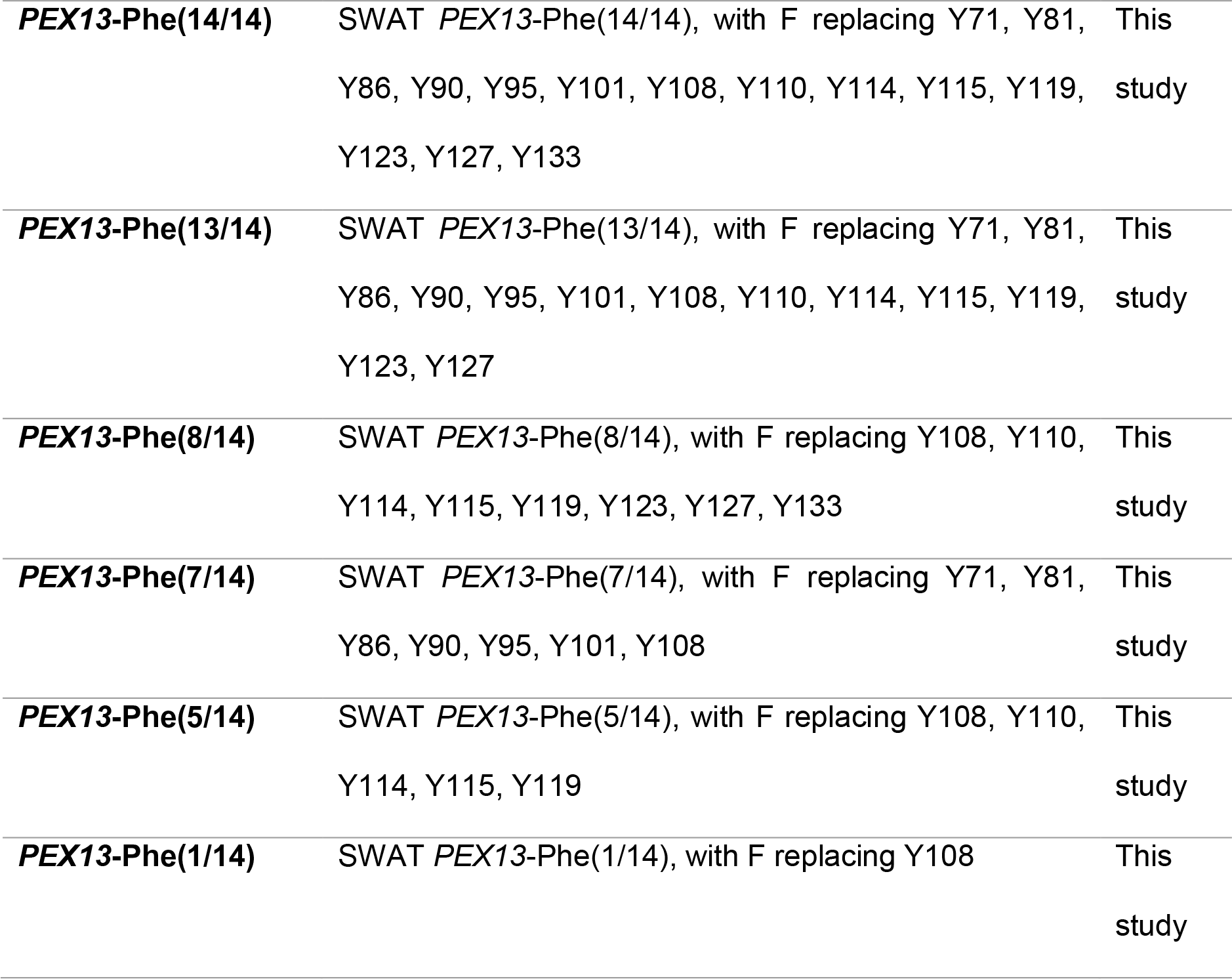
Strains used in this study. All the strains were additionally transformed with p413 ADH1 mCherry-SKL.

**Table S2.**
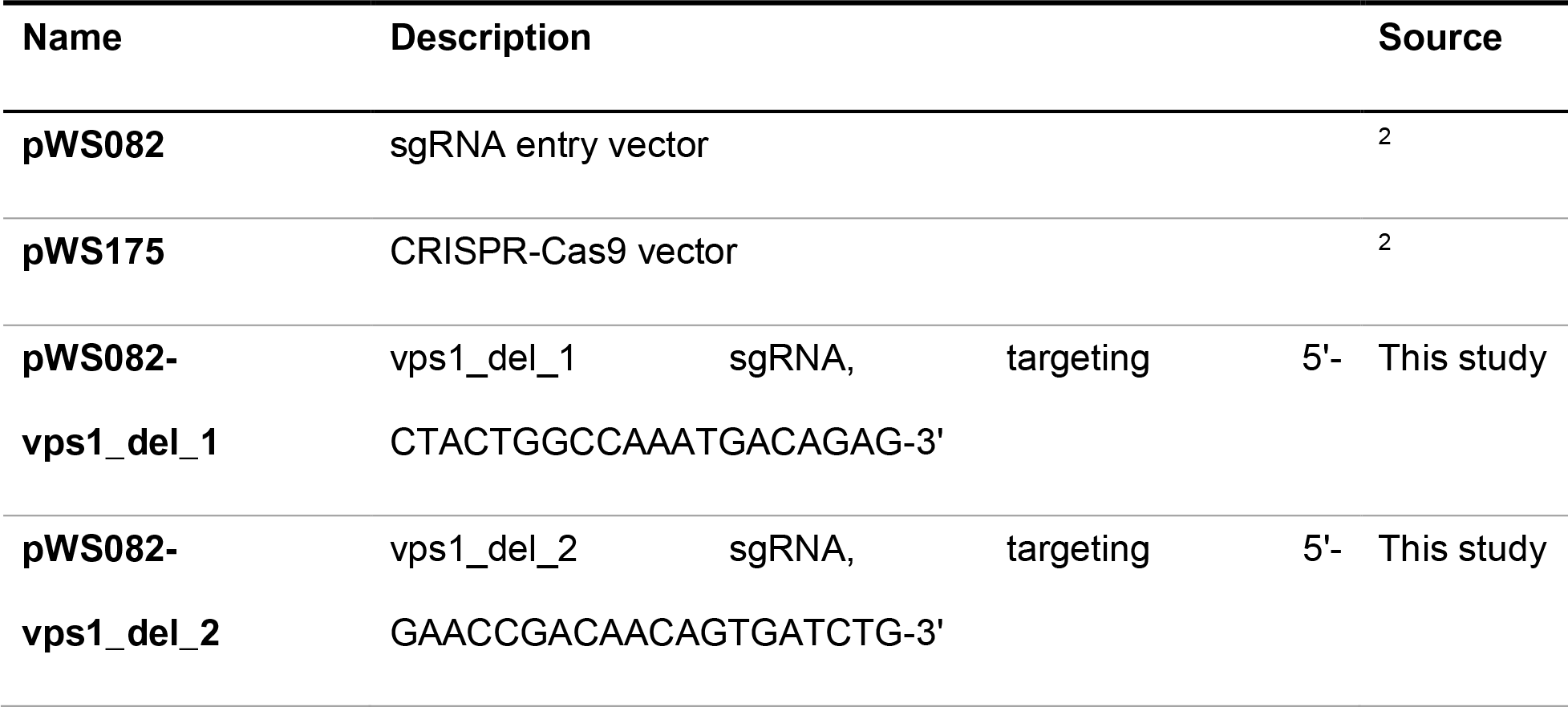
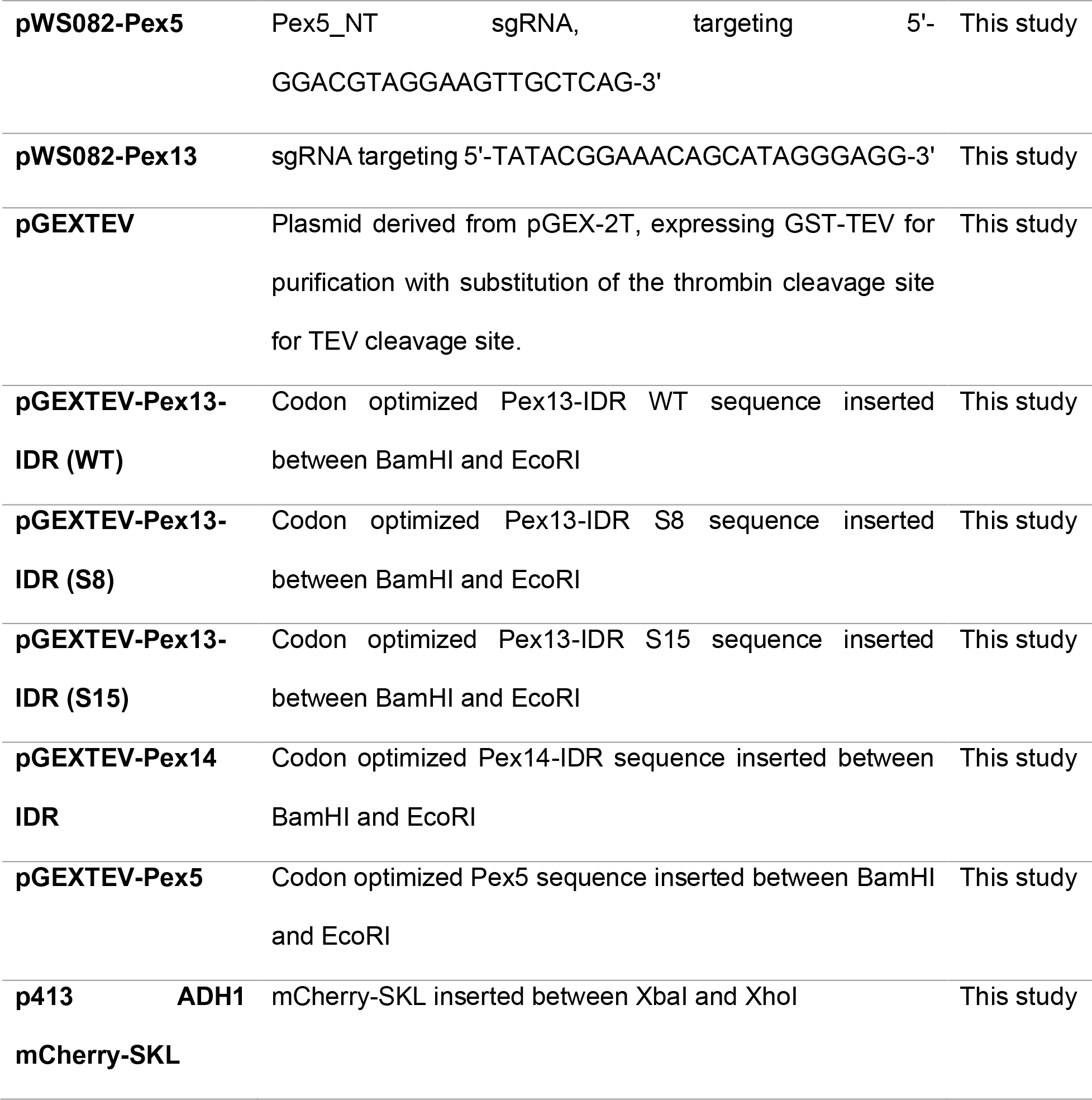
Plasmids used in this study.

